# FAVABEAN and FALAPhyl: Open-Source Pipelines for Scalable 16s rRNA Microbiome Data Processing and Visualization

**DOI:** 10.1101/2025.08.19.670115

**Authors:** Afnan Faridoun, Ramon Carvalho, Jacob Smith, Andrew Gibb, Lavanya Jain, Annie Zhang, Ajay Sran, Johanna Redmond, Mohammad Zubbair Malik, Monica Gibson, Anum Haider, Umar Rekhi, Anjali Bhagirath, Leigha D. Rock, Khaled Altabtbaei

**Affiliations:** Department of General Dental Practice. College of Dentistry. Kuwait University; School of dentistry. Faculty of Medicine and Dentistry. University of Alberta; Department of Translational Research, Dasman Diabetes Institute. Dasman. Kuwait city. Kuwait; High School Youth Researcher Summer (HYRS) Program, University of Alberta; Department of Periodontology, School of Dentistry. Indianapolis, Indiana, USA; Faculty of Dentistry, Dalhousie University, Halifax, Canada; Department of Pathology, Faculty of Medicine, Dalhousie University, Halifax, Canada; Department of Anatomical Pathology, QEII Hospital, Nova Scotia Health, Halifax, Canada

## Abstract

Reproducible and scalable analysis of 16S rRNA amplicon sequencing data remains a persistent challenge in microbiome research due to the complexity of available tools, incompatibilities between platforms, and the need for extensive bioinformatics expertise. We developed two containerized workflows—FAVABEAN (Fast Amplicon Variant Annotation, Binning, Error-correction And ANalysis) and FALAPhyl (Forays into Automating Laborious Analyses of Phylogeny)—to address these challenges. FAVABEAN and FALAPhyl are Snakemake-based pipelines designed for flexible execution across local, cluster, and cloud environments. FAVABEAN automates preprocessing, ASV inference, and taxonomic assignment using DADA2 and FIGARO, including primer averaging when samples are sequenced with multiple primers. FALAPhyl supports downstream analysis including alpha/beta diversity, network analysis, and differential abundance testing, with integrated provenance tracking. We validated both pipelines using three case studies involving oral microbiome datasets. In Case Study 1, we compared oral microbiota across family members and niches, showing primer-dependent variability in ASV-based similarity and minimal reseeding from familial sources after prophylaxis. Case Study 2 analyzed dental aerosol samples, revealing no significant microbial differences between pre-, intra-, and post-procedure air. Case Study 3, a randomized trial of a nitrate mouthrinse, demonstrated no significant microbiome shifts, highlighting oral microbial stability. FALAPhyl’s integration of DAtest enabled empirical evaluation of multiple statistical tests, aiding robust differential abundance inference. FAVABEAN and FALAPhyl offer a reproducible, automated solution for 16S rRNA amplicon data analysis. Their modular design, containerization, and provenance tracking enhance accessibility and scientific rigor in microbiome research.

## Introduction

The past decade has seen an explosion in the availability and complexity of bioinformatics tools designed to analyze microbiome sequencing data. These tools have transformed how researchers interrogate microbial communities. However, with this growth has come increased fragmentation—different tools are optimized for different data types, require incompatible file formats, or demand a deep understanding of each tool’s assumptions and parameters. This has made reproducibility and provenance tracking a persistent challenge in microbiome research(1).

One major step toward addressing these challenges was the development of QIIME2(2), a framework that emphasized transparency and data provenance. QIIME2 standardized data formats and analysis steps, making it easier to reproduce and audit workflows. However, this modular ecosystem comes with its own limitations: data must often be exported or reformatted for use in other tools, and the user must still know which steps to connect and how to parameterize them. Additionally, certain widely-used packages —such as DADA2(3) — offer simplified and less flexible implementations within QIIME2 compared to their stand-alone versions, potentially limiting analytical resolution or customization.

Other workflow management systems include mothur(4), Galaxy(5) and nf-core/ampliseq(6) provide varying degrees of automation, scalability and reproducibility but are either difficult to customize, dependent on specific platforms or require a steep learning curves for the new users.

To address these gaps, we developed and validated two complementary, open-source pipelines-**FAVABEAN** (*F*ast *A*mplicon *V*ariant *A*nnotation, *B*inning, *E*rror-correction *A*nd *A*Nalysis) and **FALAPhyl** (*F*orays into *A*utomating *L*aborious *A*nalyses of *Phyl*ogeny). These pipelines are specifically tailored for reproducible, automated analysis of 16s rRNA amplicon data enabling rapid and scalable workflows with minimal intervention. The impetus of the pipeline is to enable rapid generation of results suitable for exploratory data analysis with minimal to no post-execution hands-on interventions. The two open-source pipelines are built on Snakemake(7), making them highly scalable across different computational environments.

FAVABEAN focuses on preprocessing, denoising and taxonomic assignments of amplicon data with integrated primer-averaging functionality for datasets sequenced using multiple primer sets (8). FALAPhyl, is designed for downstream analysis and visualization, including alpha/beta diversity, network analysis, and differential abundance testing, while maintaining full data provenance via containerization and workflow tracking. Both pipelines are implemented using Snakemake, a scalable workflow management system and utilize several open-source packages, accessible under different platforms such as R, Python, and the shell(9–19). Importantly, they are containerized via Docker, ensuring bversion control and computational reproducibility across platforms-from local machines to cloud computing environments.

In the following sections, we demonstrate the functionality and capabilities of these pipelines using three diverse case studies, based on original and secondary analyses of publicly available datasets. We highlight how both FAVABEAN and FALAPhyl streamline complex analyses, reduce manual effort, and improve reproducibility— making them valuable tools for microbiome researchers across disciplines. The code for both pipelines are available on github (github.com/khalidtab/) and are available on Docker Hub as precompiled images (hub.docker.com/repositories/khalidtab).

## Materials and methods

Detailed explanation for both pipelines is available in supplementary. FAVABEAN and FALAPhyl are implemented using Snakemake(7) as its foundation, with various connecting scripts written in R, Python, and shell [Figure 1]. Snakemake itself allows the pipelines to be implemented in various scales from a single computer to a cluster, to a cloud implementation without needing to change the workflow. Although the precompiled pipelines are available as a Docker image which are pinned to specific versions of the libraries that are Linux compatible, re-compiling the pipelines dependencies to other operating systems are possible through removing the pinned files in the workflow/envs/ folder. The folder also contains the versions of the packages that the pipelines are designed to work with.

**Figure 1:**
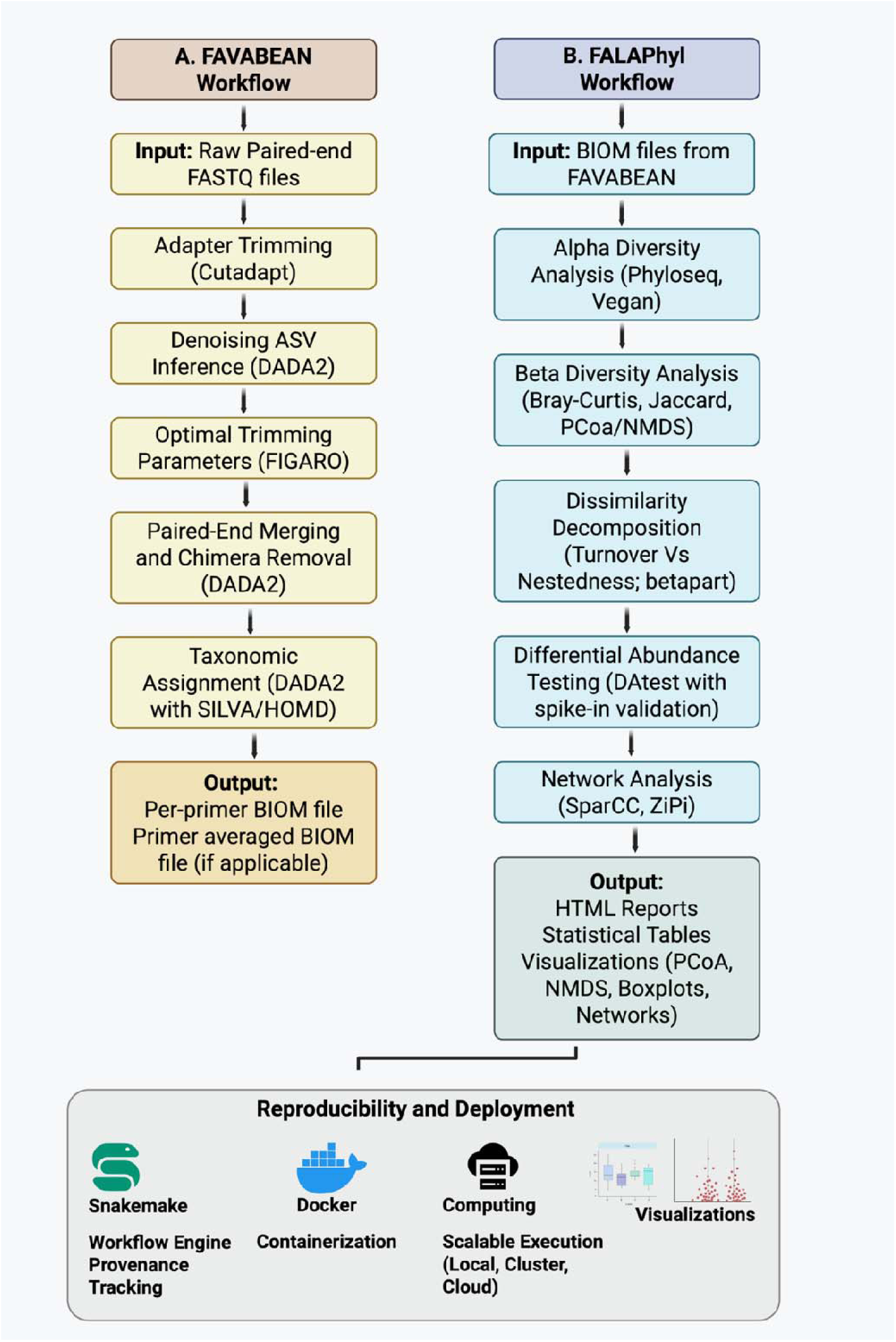
Overview of the FAVABEAN and FALAPhyl Workflow for 16S rRNA Amplicon Analysis

Within both pipelines, running the start command provides all the functions that are available to run the pipelines. Running this command also writes the requisite files for running the two pipelines; the metadata are provided as text files, and parameters are supplied to the pipelines as YAML configuration files. Description of each parameter in the YAML files are extensively explained within the files. Samples are provided in metadata text files, and parameters and metadata are specified in the pipeline the FAVABEAN pipeline, taxonomic binning reference libraries are automatically downloaded if they have not been downloaded previously.

### FAVABEAN

The pipeline’s input is FASTQ files. The files are analyzed with FIGARO(20) for the best parameters to funnel to DADA2(3) for generating Amplicon Sequence Variants (ASVs). The pipeline supports two options funnels from FIGARO; “highest coverage” which retains the greatest number of sequences in the samples at the expense of more sequencing errors analyzed by DADA2, or “lowest errors” which retains only the highest quality sequences at the expense of reducing the coverage within the samples. If two primers are used to sequence the same samples for larger coverage of the taxonomy(8), then the pipeline automatically generates a primer averaged BIOM file, in addition to the per-primer BIOM file. These BIOM files can then be used in the second pipeline to generate preliminary analyses.

### FALAPhyl

The entry points for the pipeline is through requesting the following general analyses: alpha-diversity, beta-diversity, differential abundance, network topography, and breakdown of Bray-Curtis and Jaccard dissimilarities into its two components(19). Moreover, the pipelines provide within-sample versions of alpha-diversity, differential abundances (DA), and breakdown of the two dissimilarity matrices. By utilizing Snakemake’s reporting capabilities, provenance is maintained by packaging the input files, the results, and the environments by which the results are generated into a file that can be shared to other researchers or attached to manuscripts.

The results will demonstrate the usage of these pipelines with three diverse case studies to show how the pipelines can be utilized.

## Results

### Case study 1: Microbial similarity in family members

To illustrate the functionality utility of the two pipelines, we analyzed our previously published dataset (PRJNA1159177). Detailed methods are provided in **Appendix**. In summary, oral samples from parents and their children were collected across multiple oral niches to examine microbial similarity and niche-specific differences within family units. One child underwent dental prophylaxis and was re-sampled one-week post-treatment. Samples were sequenced using two primer sets, V1-V3 (27F-519R) and V4-V5 (515bF-926R), which differ in taxonomic coverage and bias towards certain bacterial species (8). Given that differences were previously examined at a species level in the Human Microbiome Project(21), we were interested in identifying these differences at the ASV level, as well as the species taxonomy level. FAVABEAN pipeline was executed using the “pairedtaxonomy” workflow after accounting for the sample structures in a mapping file (see appendix for details and tutorial on how to run the pipeline). This generated three count tables in BIOM format: one for each primer, as well as the primer-averaged taxonomy table at the species level. The BIOM files were then processed through FALAPhyl for alpha and beta diversity analyses using both cross-sectional and paired-sample modes. Beta diversity analysis (PhILR distance) revealed distinct clustering of supragingival and subgingival plaque samples from other oral niches across all three count tables (**Figure 2**).

**Figure 2:**
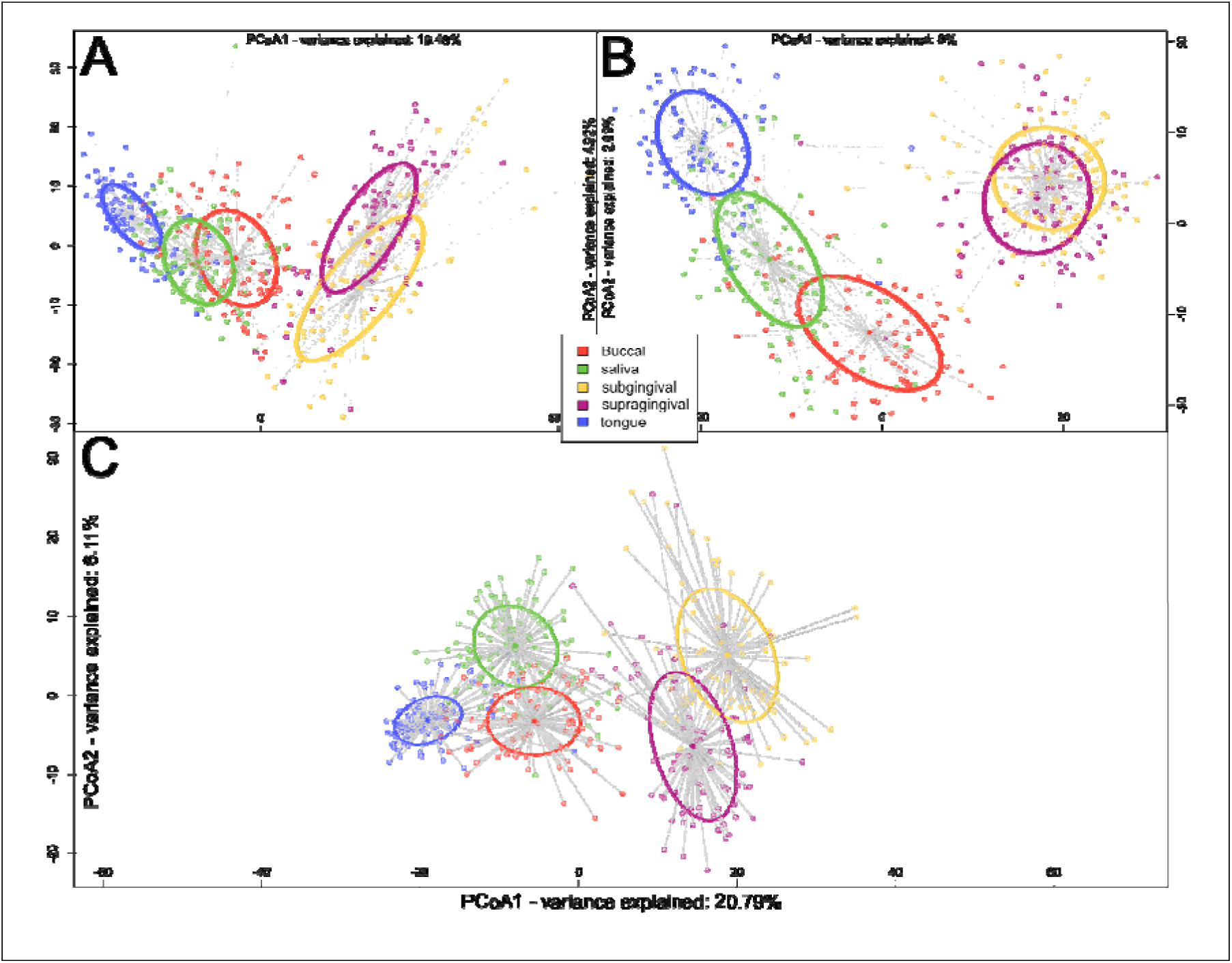
Beta diversity (PhILR distances) plots for A) V1–V3, B) V4–V5, and C) primer-averaged taxonomies. Graphs were adjusted for clarity by modifying axis labels into the graphs and legend placement.

To evaluate how primer choice affects observed microbial similarities across niches, we compared ASV-level analyses by individual primers with primer-averaged taxonomy-based results. In particular, we focused on identifying which biological sample type is most similar to subgingival and supragingival plaque. Primer-averaged taxonomy analysis showed the highest Z-scores from Dunn’s test when comparing subgingival and supragingival plaque to buccal mucosa, and lowest when comparing plaque to saliva (Z = 39.8 for buccal vs. 20.0 for saliva; both FDR-adjusted *p* < 0.05). This suggests that, among the three sampled surfaces, saliva is the most similar niche to plaque in taxonomic composition. However, when examining ASV-level differences by primer, the patterns diverged. With the 27F primer, tongue samples were the most similar niche to plaque (lowest Z-score = 5.82), while buccal mucosa showed the greatest separation (Z = 7.02); though both differences were statistically significant (FDR-adjusted *p* < 0.05), the overall distinctions were more modest than those seen in the taxonomy-based analysis. In contrast, with the V35 primer, the pattern was reversed: buccal samples were closest to plaque (Z = −2.06), while saliva was the most dissimilar (Z = −5.72). These findings highlight that conclusions about biological similarity can vary significantly depending on the primer used and the resolution of analysis (taxonomy vs. ASV). Based on these findings, the V1–V3 primer set was selected for further analysis.

Next, we analyzed the within-subject microbial dissimilarities across various oral niches to examine site-specific variations. FALAPhyl was run again on a new mapping file containing only paired samples of saliva, sub- and supra-gingival plaque at the baseline time point. The pipeline itself automatically excludes unwanted samples and runs the analysis without needing to pre-filter the biom count table. Paired-sample analysis (paired-alpha paired-beta paired-diff) was done. The observed ASV richness between supragingival and subgingival plaque was not significantly different (Wilcoxon signed-rank test, p > 0.05; Figure 3B).

**Figure 3A:**
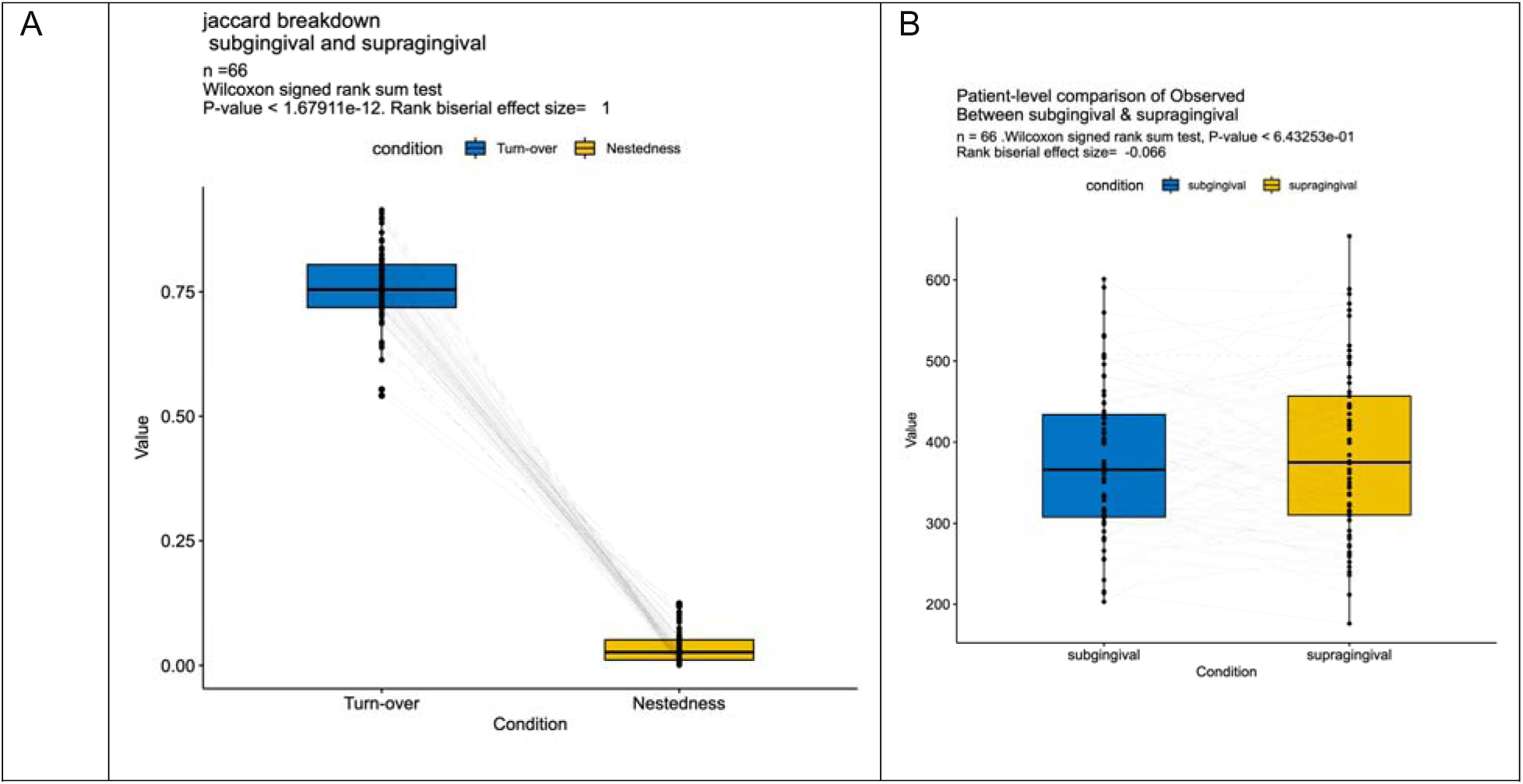
Dissimilarity between supragingival and subgingival plaque in microbial membership within the same participant, illustrated through the two components that form the Jaccard dissimilarity; turnover, and nestedness. Figure 3B: Observed features alpha diversity between supragingival and subgingival plaque samples within the same individual.

Given that supragingival plaque is continuous with subgingival plaque, and is more readily exposed to saliva, we were interested in understanding how similar it is to saliva compared to subgingival plaque. To illustrate the ASV membership dissimilarity between sites, we used Jaccard dissimilarity. Jaccard dissimilarity analysis showed that saliva was equally dissimilar to both supragingival and subgingival plaque (Friedman’s test, FDR-adjusted p = 0.3). Therefore, we wanted to focus on the differences between supra- and sub-gingival plaque. Jaccard dissimilarity was broken down to its two components – turnover (replacement of an ASV by another) and nestedness (non-replacement leading to one site being a subset of the other)(19). Dissimilarity decomposition revealed significantly greater turnover than nestedness between supragingival and subgingival plaque (Wilcoxon signed-rank test, p < 0.05; Figure 3A).This indicates that the missing ASV in one site is replaced with one not found in the other site, so much so that the number of observed ASVs in the two was not significantly different from each other (p>0.05 – 05).

By utilizing the internal reporting capabilities of Snakemake (snakemake include_biom_and_meta alpha beta breakdown diff network paired_beta paired_alpha paired_diff --report data/report.html), the original files, steps taken to generate the results, and the steps needed to generate them are packaged into a zipped file, thereby enabling provenance and transparency without locking the files into a specific format. This step also prevents erroneous attribution of analyses to incorrect pipeline runs (eg: full samples vs only paired samples in this example) but packaging only the correct results in the report. This method also allows for custom analysis to be done as we have below, without needing to extract artefacts as is done with QIIME2.

We were interested in identifying the percentage of ASVs that were present in the one-week samples in the supra- and subgingival plaque samples that were not present in the child the week before, but were found in other family members, as well as those not linked to any familial sources. Reported as mean±standard deviation, the percentage of ASVs attributable to familial non-child sources was 0.169%±0.08%, which was smaller than those not attributable to any familial sources (0.548%±0.149%). However, the familial sources, represents a two-fold difference in the mean relative abundances compared to non-familial sources, (0.0206%±0.0126% and 0.0125%±0.0044%, respectively).

### Case study 2: Fallow time in dental clinical settings

In our previously published study(22), we analyzed the bacterial content in dental aerosols before, during and after procedures to identify potential sources and evaluate risks. In this secondary analysis, we compared post-procedure air samples to pre- and intra-procedure air to assess microbial dissimilarity and inform infection control practices (Figure 4A–B). Our goal was to determine whether the microbial composition of post-treatment air differed significantly from the baseline to assess the necessity of fallow time in dental settings.

**Figure 4:**
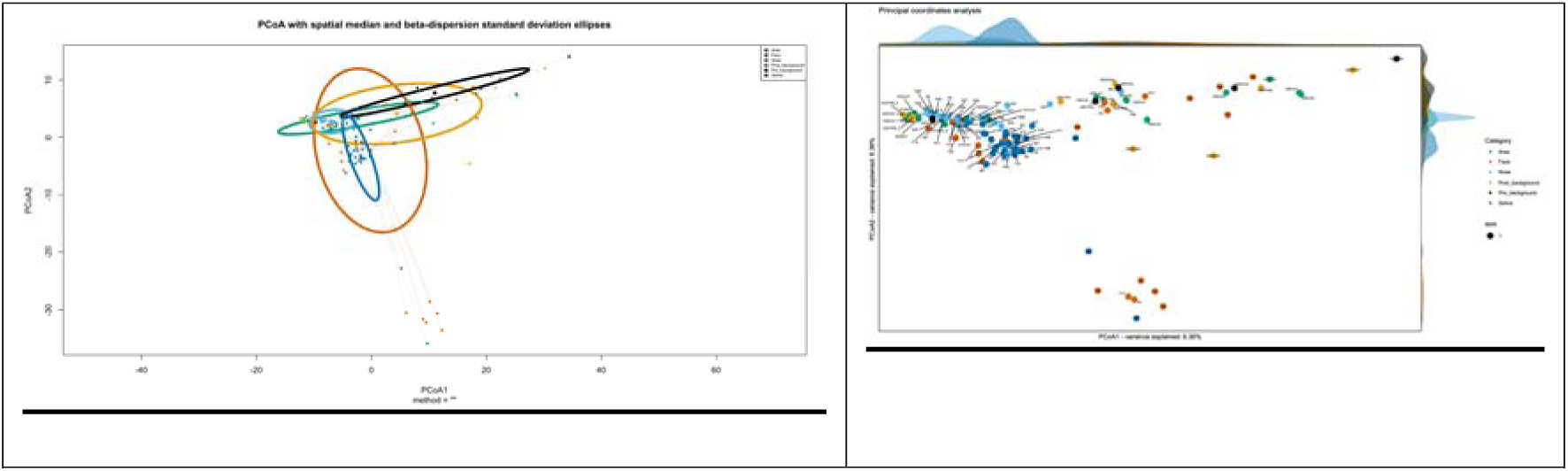
Principal Coordinates Analysis (PCoA) based on PhILR distances.(A) PCoA plots show dispersion using the standard deviation ellipses (representing one standard deviation of the spread of the samples away from the centroid, which may become cumbersome to illustrate any differences). (B) An alternate visualization of the same data using the probability density functions along principal axes, providing clearer distributional insight. The pipeline also generates Sample-labeled PCoA plots automatically.

Beta diversity analysis (group centroids, ADONIS) revealed no significant differences between post-procedural and pre-procedural air (ADONIS, FDR-adjusted p = 0.578; ANOSIM, FDR-adjusted p = 0.159), nor between post- and intra-procedural air (ADONIS, p = 0.49; ANOSIM, p = 0.32).

### Case study 3: Nitrate mouthrinse and microbiome stability

A parallel arm randomized clinical trial was conducted to test the effect of a nitrate-rich mouthrinses on the oral microbiome composition (details in **Appendix 3**). In total, 21 participants were randomized to either the intervention group (n = 11; nitrate mouthrinse) or the placebo control group (n = 10) administered daily for 14 days. To enable species-level resolution, primer-averaged BIOM outputs from FAVABEAN were used for downstream exploratory analysis. No significant differences in overall microbial composition were observed between intervention and placebo groups (ADONIS and ANOSIM; p > 0.05 for both tests). We next explored whether subtle species-level differences could be detected despite the absence of global shifts. Given the lack of a universally accepted method for differential abundance analysis — and the variability of results across methods — we employed the DAtest package to systematically assess test performance on this dataset. DAtest package uses artificial spike-in simulations to empirically evaluate statistical test performance under varying conditions. Our pipeline integrates this approach to guide method selection based on dataset-specific resilience. Our modifications of the DAtest package (parallelization to the spiking tests, and custom graphing outputs with incorporation of an additional test) are detailed in **Appendix 3**. None of the paired statistical testing identified significantly differentially abundant species in either the intervention or the control groups (FDR adjusted p>0.05 for all tests). Moreover, none of the methods had a score above zero, indicating that none of them are able to reliably identify the statistically significant features (Figure 5).

**Figure 5:**
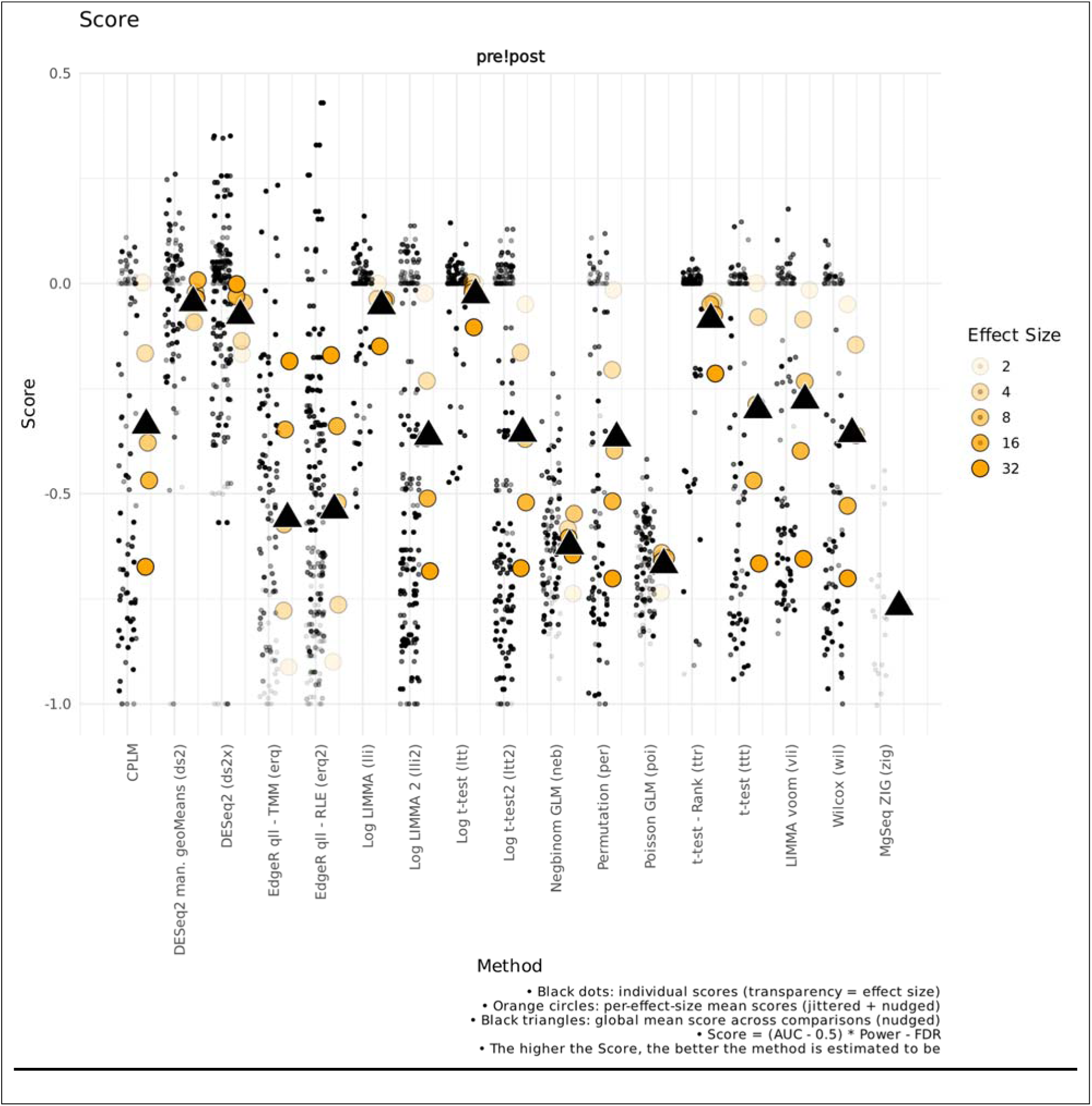
Scores from the paired testing in Intervention arm, baseline and post-intervention paired samples. The test shows the dataset interrogated against multiple spike-in trials, with different effect sizes. Black triangle illustrates the average of the effect sizes. As none of the averages reach the zero threshold, none of the methods are capable of reliably identifying the statistically significant features.

## Discussion

### Technical Insights

FAVABEAN and FALAPhyl were developed to address key challenges in 16S rRNA amplicon data analysis including the need for automation, reproducibility and scalability across diverse computing environments. Although FAVABEAN is solely designed for amplicon-based sequencing workflows in mind, FALAPhyl’s analytical framework is data-agnostic and applicable to any compositional dataset, including those beyond 16S rRNA sequencing. The breadth of analytical tools created for FALAPhyl allow for both cross-sectional and paired study designs. This enables users to efficiently and comprehensively perform exploratory and hypothesis-generating analyses with minimal manual intervention. Together, FABABEAN and FALAPhyl demonstrate how containerization and workflow-based approaches can streamline complex bioinformatics analyses while maintaining transparency, version control and scientific rigor. The integration of Snakemake as the workflow engine for both pipelines provide several key advantages over traditional script-based analyses including scalability, reproducibility and automated provenance tracking. The automatic scaling from single computers to cloud implementations without the need for specialized infrastructure removes technical barriers that often limit researchers’ ability to process large datasets. By containerizing both pipelines, consistency across different computational environments eliminating issues related to software version discrepancies-a common source of irreproducibility in microbiome research(1). FALAPhyl’s integration of the DAtest framework addresses a critical challenge in the selection of a suitable differential abundance methods in the absence of a gold-standard approach. By empirically testing multiple approaches on each dataset through spike-in simulations, DAtest enables researchers to select the most robust statistical approaches based on their performance with the specific data characteristics rather than relying on general recommendations. We enhanced DAtest within FALAPhyl by parallelizing spike-in simulations, which improves computational efficiency and enables its application to larger datasets. Moreover, the FALAPhyl’s automated handling of paired samples is particularly advantageous for longitudinal and intervention studies, where manual subsetting is error-prone and labor-intensive. Traditional approaches often require manual data subsetting and careful tracking of sample relationships, introducing opportunities for error. Automating these workflows reduces analytical burden and potential for user error, while Snakemake’s integrated reporting ensures complete provenance and reproducibility.

### Biological Insights from Case Studies

#### In Case study 1, we assessed microbial similarity across family members and oral niches

<u>While</u> second-generation sequencing has massively lowered the cost of DNA sequencing compared to that of Sanger sequencing, It sacrifices sequence read-length to compensate for parallelization of sequencing. This serves as a challenge; Sanger sequencing can sequence the much longer stretches of the 16s rDNA gene, which provides excellent identification capability of the taxonomy through coverage of the regions that contain the characteristic mutations of these taxa. This was not the case for second-generation sequencing technologies such as Illumina sequencing. Therefore, alternate strategies, such primer-averaging, were employed to provide the similar breadth of taxonomic coverage and sample composition. The development of ASVs allowed for more granular analysis at the genome level, which does not allow for primer averaging to occur. Our results demonstrate that ASV-level and taxonomy-based analyses yield different biological interpretations, underscoring the importance of analytical consistency and caution when comparing across various levels of resolution.

Microscopic examinations have shown the supra and sub gingival plaque are anatomically contiguous, allowing for continuous microbial exchange between these niches. The gingival crevicular fluid carries planktonic-state bacteria, as well as flecks of biofilm from the subgingival environment toward the oral cavity, thereby meeting the supragingival plaque. This provides avenues of microbial dispersal between the two environments. Dissimilarity decomposition revealed that inter-site variation was driven primarily by species turnover rather than nestedness, suggesting that each site harbors distinct microbial communities adapted to its environment. Therefore, one should not assume that ecological filtering by way of changes in the environment such as reduced Oxygen tension, or changes in the nutritional resources in the environment, caused attrition in the microbial composition resulting in their differences. Rather, the driver of the differences between the two is largely due to the distinct membership in the two sites, likely adapted to exploit the ecological differences in the two sites.

The minimal colonization by family-sourced ASVs following prophylaxis (0.169% ± 0.08%) provides quantitative support for the stability and resilience of individual oral microbiomes. While family members share environmental exposures, the re-establishment of the oral microbiome appears to be driven predominantly by the individual’s own microbial reservoir rather than horizontal transmission from family members. Although is not a novel finding onto itself, the novelty is in the fact that such seeding is detectable at all. To our knowledge, this is the first study to quantify the contribution of non-host familial microbiomes to recolonization of the subgingival plaque niche. The majority of previous studies have concentrated on maternal/child relationships, rather than whole family transmissions(23). As similarity in the oral microbiota between the mother-child dyad increases during childhood and adolescence(23), our results provide quantitative evidence of such seeding. Oral prophylaxis is used to remove the supragingival plaque of the child as mean to “professionally” brush teeth. Supragingival plaque debridement, and especially professional debridement, has long been known to have an effect on the subgingival clinical status, and its microbiome(23–31), with the effect likely negatively associated with the increasing depth of the periodontal pocket. This is the first time that seeding from outside sources of the person’s the oral cavity be attributed to the contribution of the restitution of the subgingival microbiome. While we followed the participants for only one week, it is possible that further maturation of the biofilm would reduce the external contributions as the resident microbiota regain control over their niches. We suspect this is only partially true as dispersion events occur daily through tooth brushing, which has been shown to enhance the microbial similarity between twins(32). Therefore, we believe regular oral hygiene likely increases the acquisition from external sources, through a process of repeated dispersal events. Familial ASV transmission was approximately twice that of non-familial sources, warranting further investigation into close-contact microbial exchange within households.

In this case study, the primerl1laveraging module of FAVABEAN enabled seamless consolidation of V1–V3 and V4–V5 amplicon data, reducing primer-driven bias in downstream taxonomic inference. Simultaneously, FALAPhyl’s automatic pairedl1lsample analysis allowed within-subject beta diversity and differential abundance testing without preprocessing steps, preserving analysis provenance and minimizing oversight. These integrated features facilitated clear, reproducible assessments of family microbial similarity that would be cumbersome in manual multi-primer workflows.

#### In Case study 2 we assessed the need for fallow time in dental clinical settings

Fallow time, introduced during the COVID-19 pandemic, represented a logistical challenge by delaying post-patient practice turnover to allow for aerosol particles to settle to the floor. Our analysis showed that microbial composition in post-procedural air was statistically indistinguishable from pre-procedural air samples. This finding has practical implications for infection control protocols in dental practice. Aerosolized bacteria either settle or disperse rapidly, potentially reducing the need for extended fallow times between patients, or that relying on the bacterial content of the air from known sources as a surrogate to air contamination is not a good surrogate to actual quantitative measurement of the microbial agent in the air. Interpretation of these findings warrants caution. Larger studies incorporating quantitative pathogen detection methods (e.g., qPCR or culture-based assays) are needed to validate microbial risk assessments based solely on amplicon sequencing data.

In this case study, FALAPhyl’s automated beta diversity generation and paired-group testing demonstrated that post-procedural air microbial composition was statistically indistinct from pre-procedural levels. These detailed comparisons — powered by Snakemake-driven provenance, PhILR distance calculations, and ADONIS/ANOSIM tests — provide timely, reproducible evidence relevant to infection control decisions, and could be replicated or extended with minimal reconfiguration.

#### In case study 3 we assessed the efficacy of a nitrate-rich mouthrinse on oral microbiome stability

The absence of detectable microbiome shifts following the use of the nitrate mouthrinse highlights the resilience of oral microbiome. Despite nitrate’s known role in promoting nitrate-reducing bacteria and potentially influencing oral and systemic health through the nitrate-nitrite-nitric oxide pathway, we observed no compositional shifts after 14 days of daily use. This stability likely reflects several biological factors: the tongue microbiome’s inherent resilience in healthy individuals(21), the possible inadequacy of once-daily dosing to exert sustained selective pressure, and the short intervention period relative to the stability of established microbial communities. These findings align with emerging evidence that healthy oral microbiomes resist perturbation from single-agent interventions(33). Unlike the disruption seen with broad-spectrum antimicrobials, a metabolic substrate like nitrate may require either longer exposure, higher frequency of use, or a dysbiotic baseline state to induce measurable shifts in the community composition. This is consistent with the *Anna Karenina’s principle* where dysbiotic microbiomes exhibit greater variability compared to stable, more homogenous healthy profiles(34). The participants’ healthy periodontal likely contributed to this observed microbial stability, as diverse and balanced communities typically demonstrate greater resilience to external perturbations(33). These results suggest that microbiome-targeted interventions in healthy populations may need to consider the ecological stability of established communities. Future studies should consider recruiting individuals with existing dysbiosis and extending intervention duration to evaluate whether metabolic shifts precede detectable compositional changes.

This case study highlights how FALAPhyl incorporated DAtest-based empirical evaluation of several differential abundance methods using datasetl1lspecific spike-in simulations. This guided selection of appropriate tests and validated the null finding of no significant microbiome shift, reinforcing confidence in the result’s validity. Without this framework, researchers would lack both systematic test selection and documentation of method suitability.

### Comparison with Existing OpenlZISource Amplicon Analysis Pipelines

Recent years have seen the widespread adoption of several open-source workflows designed for reproducible 16S rRNA amplicon sequencing analysis, including QIIME-2 (with DADA2), nf-core/ampliseq, Tourmaline, and mothur-based pipelines. QIIME-2 combined with the DADA2 plugin achieves superior accuracy in taxonomic classification, reduced false-positive detection, and improved diversity estimation compared to earlier tools such as QIIME-1, mothur, and MEGAN.

The nf-core/ampliseq pipeline, implemented in Nextflow and distributed via Docker, extends QIIME-2 and DADA2 into a scalable, containerized workflow. It supports a wide range of sequencing platforms (Illumina, PacBio, IonTorrent) and marker types (16S, ITS, COI), generating MultiQC summaries and interactive QIIME-2 outputs. Similarly, Tourmaline integrates Snakemake and QIIME-2, offering automated execution and reporting within a reproducible Snakemake framework.

Relative to these tools, FAVABEAN and FALAPhyl provide complementary and distinct features. Unlike other pipelines, FAVABEAN automatically averages taxonomic outputs across multiple primer sets sequenced from the same sample. This minimizes primer bias and allows a more comprehensive taxonomic inference, a feature unique to this workflow. FALAPhyl integrates the DAtest framework using spike-in simulations to empirically evaluate and guide the choice among multiple statistical tests. This dataset-specific method selection is not available in standard QIIME-2 or nf-core workflows. FALAPhyl also automatically processes paired or within-subject comparisons in a single command, eliminating manual subsetting steps and reducing error potential, which is especially valuable for longitudinal or intervention studies. Built entirely on Snakemake, both pipelines support execution across local, cluster, or cloud environments without modification. Snakemake’s provenance tracking ensures complete transparency and reproducibility. Lastly, deployment via Docker ensures version control, platform independence, and consistent results across computing environments. However, in contrast to nf-core/ampliseq, FAVABEAN and FALAPhyl are command-line only, with no graphical user interface (GUI) support (e.g., Galaxy, QIIME Studio). Additionally, while nf-core workflows support diverse marker types and sequencing platforms, FAVABEAN is currently optimized specifically for paired-end Illumina 16S data. Furthermore, nf-core pipelines benefit from broader community adoption and support, whereas FAVABEAN and FALAPhyl are in earlier stages of community uptake and currently rely on GitHub-based documentation and user contributions.

In summary, FAVABEAN and FALAPhyl fill a critical niche in the microbiome analysis landscape: they offer advanced functionalities—including primer-averaging, empirical statistical method selection, and automated paired-sample analysis—within a reproducible, Snakemake-based framework. These features particularly benefit researchers seeking flexibility, statistical rigor, and reproducibility without dependence on large, monolithic platforms.

### Limitations and Future Directions

Several limitations of the current pipeline and its implementation merit discussion.

First, FAVABEAN currently supports only paired-end sequencing, which limits its applicability to studies using single-end read formats—a common approach in some targeted amplicon studies. Second, while many steps in both pipelines are automated, users are still required to specify key analysis parameters such as diversity metrics and distance measures. Although sensible defaults and detailed documentation are provided, a basic understanding of bioinformatics and microbiome statistics remains necessary to ensure optimal and accurate usage. Third, while the pipelines are well-suited for standard 16S workflows, novel or highly specialized analyses (e.g., rarefaction-free methods, phylogenetic network models) would require customization beyond the current implementation. The modular nature of Snakemake facilitates such extensions; however, doing so demands some programming proficiency.

Additionally, reliance on Docker containerization, though essential for reproducibility and version control, may pose access limitations for users operating in high-security or restricted computing environments where container technologies are not supported.

Despite these limitations, the open-source design of FAVABEAN and FALAPhyl promotes community-driven development, adaptation, and extension, enabling researchers to tailor the workflows to evolving needs and diverse experimental designs.

## Funding information

Case study 1 was supported by funded by the Oral Health for Children, Youth, and Families Fund (OHCY-02).

Case study 3 was supported by grant funding from the Dalhousie Medical Research Foundation (22-092).

## Contributions

Author 1. Initial idea of the pipeline. Testing pipeline and implementation improvements. Data analysis for case study 1-3. Manuscript preparation. Critical appraisal of the manuscript.

Author 2. Data collection of case study 2. Critical appraisal of the manuscript.

Author 3: Study design of case study 3. Data collection of case study 3. Critical appraisal of the manuscript.

Author 4-5: Study design of case study 1. Data collection of case study 1. Critical appraisal of the manuscript.

Author 6-7: Pipeline testing and implementation. Critical appraisal of the manuscript.

Author 8: Sample collection and preparation for case study 1. Critical appraisal of the manuscript.

Author 9: Pipeline implementation, consulting and improvement. Critical appraisal of the manuscript.

Author 10: Design of case study 1. Critical appraisal of manuscript.

Author 11-12: Pipeline improvements. Critical appraisal of manuscript.

Author 13: Design of case study 1. Sample collection for study 1. Manuscript preparation. Critical appraisal of the manuscript.

Author 14: Design of case study 3. Sample collection for study 3. Manuscript preparation. Critical appraisal of the manuscript.

Author 15: Initial idea of the pipeline. Testing pipeline and implementation improvements. Data analysis for case study 1-3. Manuscript preparation. Critical appraisal of the manuscript.

### Appendix 1

Detailed supplementary

#### FAVABEAN

**Part 1: study set up**

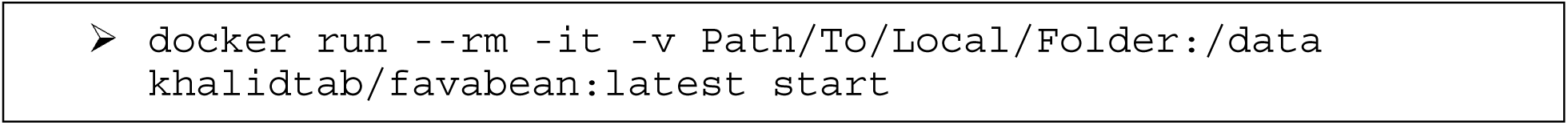

The following steps illustrate using FAVABEAN using Docker in an interactive mode.

Although the image contains both bash and sh, running the pipeline with the start will

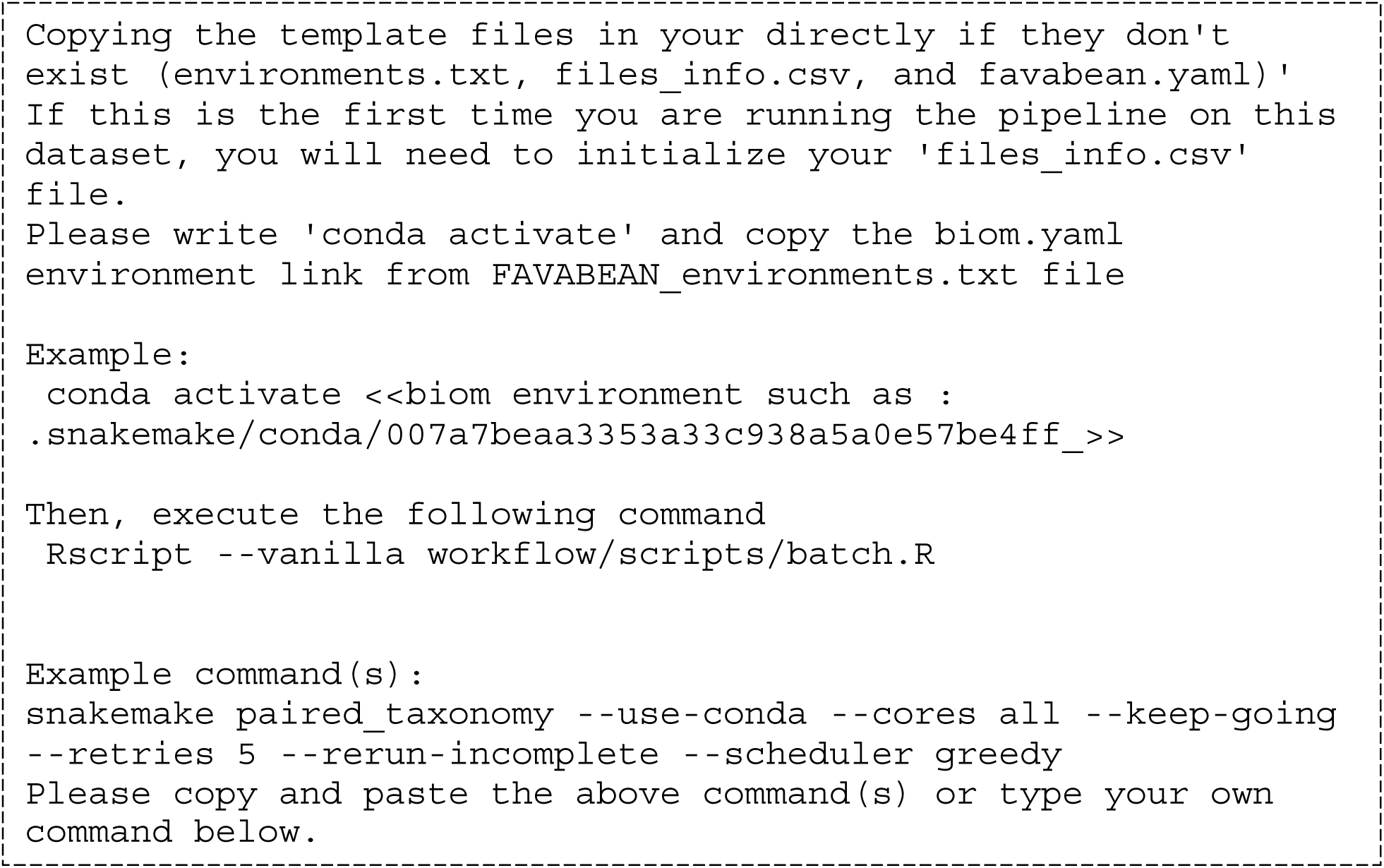

print commands needed to use the pipeline, as below:

Below is the step-by-step tutorial on using the pipeline

##### 1. Create the metadata table for the sequencing FASTQ files

As mentioned in the prompt above, when `start` was executed, files_info.csv was copied to your `Path/To/Local/Folder.` This comma-separated table is where the fastq files are identified. Below is a minimal example:

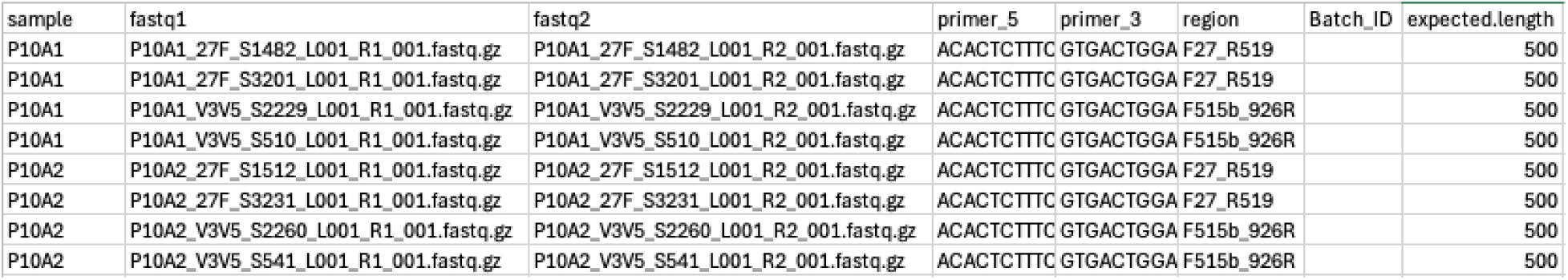

Columns:

- **sample**: This is where you note the eventual name of the samples in the biom file. Note that in this example, we have the same sample, identified by the first column, sequenced multiple times, using two different primers. The merging strategy will be discussed in later stages of this tutorial.
- **fastq1, fastq2**: path to the fastq files in your `Path/To/Local/Folder`
- **primer_5, primer_3**: the forward and reverse primers used for producing the amplicons.
- **region:** name of the sequenced region. This is used as a placeholder for the primer tables. Any name to identify the region is fine.
- **Batch_ID:** see below
- **Expected_length:** expected length of the amplicon. If unknown, the largest length is acceptable. Generally speaking, Illumina MiSeq chemistry allows for 350 bp in both forward and reverse reads. After primer sequences removal, a 500 bp length is reasonable and therefore used as default.

##### 2. (Optional) Identify sequence batches

As the machine learning algorithm Error Learning Algorithm of DADA2(3) is batch-specific, we need to identify the batch of each sample. Failure to explicitly identify the batches will likely degrade the quality of the error learning. If this information is not available, the following step is executed.

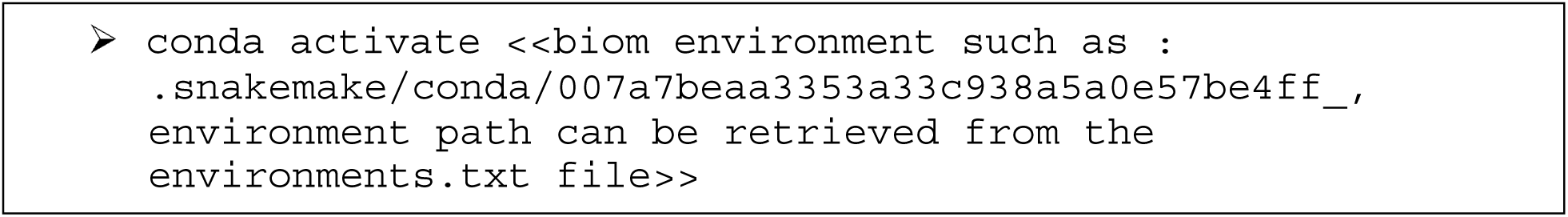

Then execute the following command

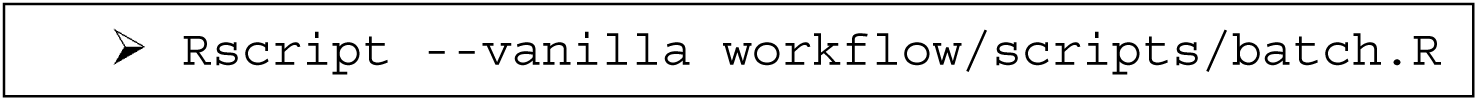

This will read the headers of each of the files that are in the Illumina CASAVA 1.8+ format, and will segment the files based on the following schema:

*@<instrument<:<run number>:<flowcell ID<:<lane<…*

Once successfully executed, a new CSV file, file_info_Batches.csv, is generated with the correct information as below:

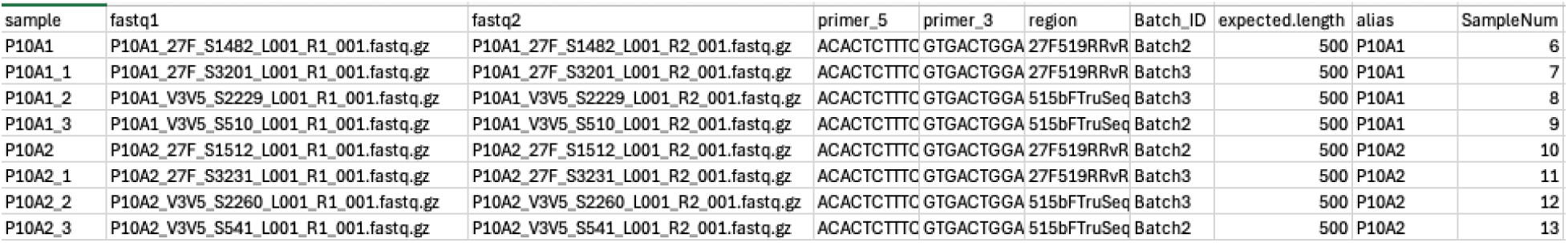

##### 3. Select the parameters for FAVABEAN

The parameters are available in the favabean.yaml file. Deselecting a parameter is simply done by commenting out the beginning of the file with “#”. Below are options:

- **initial_filter**

- This step helps remove sequences not expected to give you a meaningful hit to DADA2 because they are too short.
- **trim_param**

- Automatic identification of the trimming parameters for DADA2 typically requires manual work, which may lead to reproducibility errors. This pipeline uses Figaro for this step. Figaro expects all sequences to be of the same length, so cutting all sequences to a specific length is needed for this step. These trimmed sequences are only used in Figaro and are not used during the other steps. This trimming also simplifies the Figaro analyses, and it helps
- You can write a specific number you like (eg 100, for 100 basepairs), or, you can filter out sequences based on general parameters as below

- <u>Default</u>: is that if the length of sequences in Q1 is within 10% of Q2 (median), then use the Q1 to include the majority of sequences. Otherwise Q2 is used
- <u>Q1</u>: 25th percentile length
- <u>Q2</u>: 50th percentile length (median)
- <u>Q3</u>: 75th percentile length
- max_len: longest sequence length
- Note that this accepts one value only. So if you have more than 1 uncommented, only the first one will be used.
- **Figaro(20)**

- Figaro provides multiple options for the DADA2 parameters along with their error quality scores. The pipeline is implemented with the option of the two extremes:
- highest_coverage: this option retains the largest percentage of sequences
- lowest_errors: this option uses the parameters with the largest score.
- **taxonomy_database**

- Bacterial identification is done through DADA2. By default, the links to two databases are provided; eHOMD(35) and SILVA(36). If the path to the files is not available in your `Path/To/Local/Folder,` the files will be downloaded for the `Path/To/Local/Folder/resources`. Adding new reference databases is trivial by either identifying their downloading link in the yaml file, or by creating a placeholder entry in the yaml file, and placing the reference databases in the `resources` folder.

##### 3. Execute the pipeline

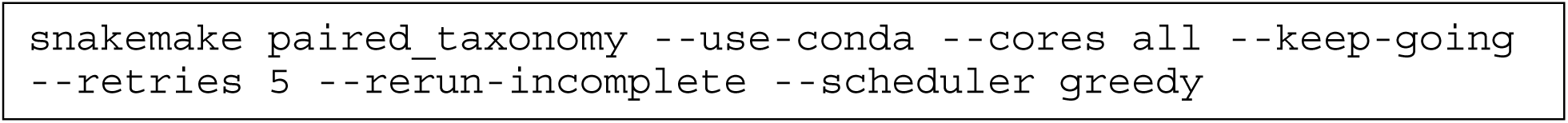

**<u>Note:</u>** Currently, only analysis of paired fastq files are implemented. Future versions will work on forward sequenced fastq files.

Two options available for analysis: `paired_taxonomy`, and `paired.` The difference between the two is that `paired_taxonomy` will identify the taxonomy of the sequences, in addition to generating the ASVs that `paired` does. Below are explanations of SnakeMake options, not specific to the pipeline.

- --use-conda: This is mandatory for the proper execution of the pipeline
- --cores: Either a numeric value is to be provided, or all for all cores available to the docker container.
- (optional) --keep-going: In case of an error, all non-dependent branches of the analysis will continue their execution.
- (optional) --rerun-incomplete: in case of an error, rerun the step
- (optional) --retries: in case of an error, retry the step X number of times.
- (optional) `--scheduler:` options are `greedy,` and `ilp.` The latter option tries to reduce runtime and disk usage by best possible use of resources, but in some cases, takes a long time to find the best path for scheduling the steps.

**Part 2: Pipeline steps explanation**

Below is the graphic representation pipeline, with explanation of each step.

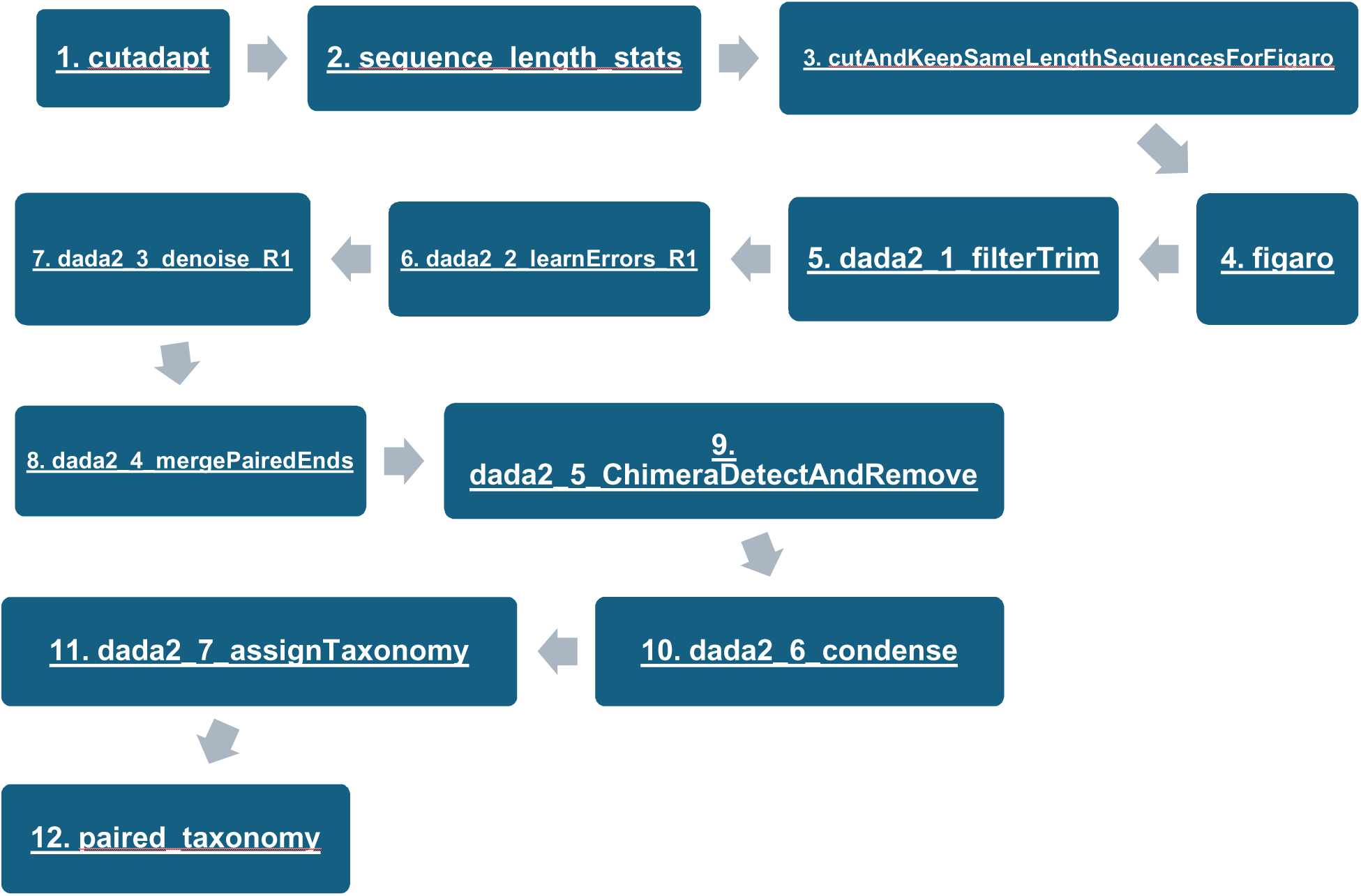

1. **<u>cutadapt</u>**

- Removes adaptors from sequences based on values in files_info_Batches.csv
2. **<u>sequence_length_stats</u>**

- Calculate descriptive statistics using the package SeqKit on measure the length of the forward and reverse sequences
3. **<u>cutAndKeepSameLengthSequencesForFigaro</u>**

- Using SeqKit, the script will filter the FASTQ files based on the descriptive statistics from the step before
4. **<u>figaro</u>**

- Calculates the optimum trimming parameters for DADA2
5. **<u>dada2_1_filterTrim</u>**

- Using the output from step 1, filtering is done based on parameters provided by step 4
6. **<u>dada2_2_learnErrors_R1</u>**

- Learns sequencing errors on forward reads
- Note: dada2_2_learnErrors_R2 is run in parallel for reverse reads
7. **<u>dada2_3_denoise_R1</u>**

- Creates the model for denoising of forward reads
- Note: dada2_3_denoise_R2 is run in parallel for reverse reads
8. **<u>dada2_4_mergePairedEnds</u>**

- Both forward and reverse denoised reads are reads and merged
9. **<u>dada2_5_ChimeraDetectAndRemove</u>**

- Chimeras are detected and removed
10. **<u>dada2_6_condense</u>**

- In-house developed faster version of <u>DADA2_COLLAPSE_NOMISMATCH</u> function. DADA2 version loops through all sequences one by one and determines whether they are “collapsable”. The optimized version precomputes the prefixes, and groups them by these prefixes prior to the loop that determines if these groups are collapsable.
11. **<u>dada2_7_assignTaxonomy</u>**

- Assigns taxonomy based on the selected database reference
12. **<u>paired_taxonomy</u>**

- If two primers exist for each sample, then primer averaging is performed

#### FALAPhyl

**Part 2: Pipeline steps explanation**

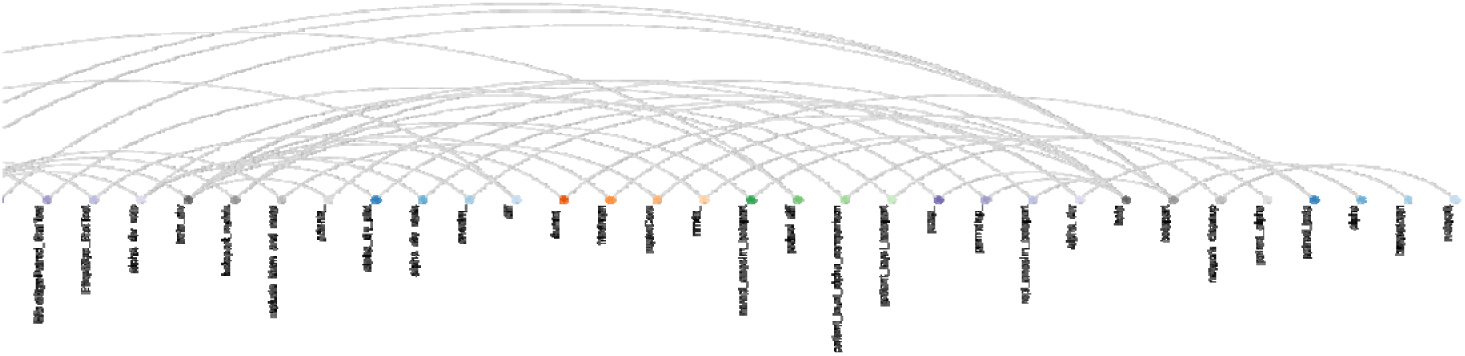

Above is the graphic representation of the steps, and their inter-connectedness. Due to its complexity, we will focus on the endpoints of the pipeline as accessible through the command line to run them all:

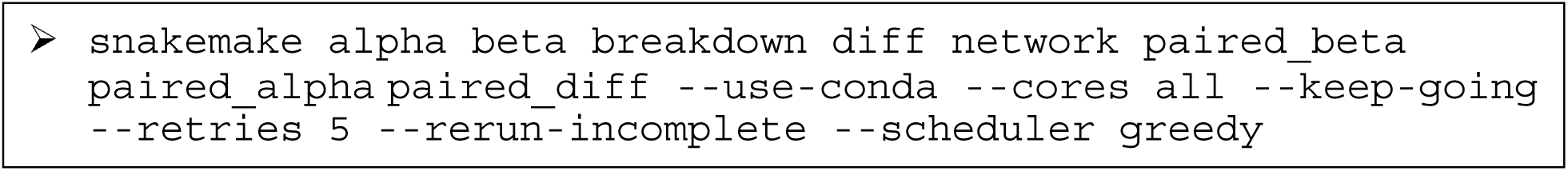

Below, any variables defined in the falaphyl.yaml file that are used in the explanation are written inside square brackets “[variable name]”. The graphs are provided as they would be generated through the algorithm using the generic sizes that are written by default in the falaphyl.yaml file, but they may be adjusted as needed.

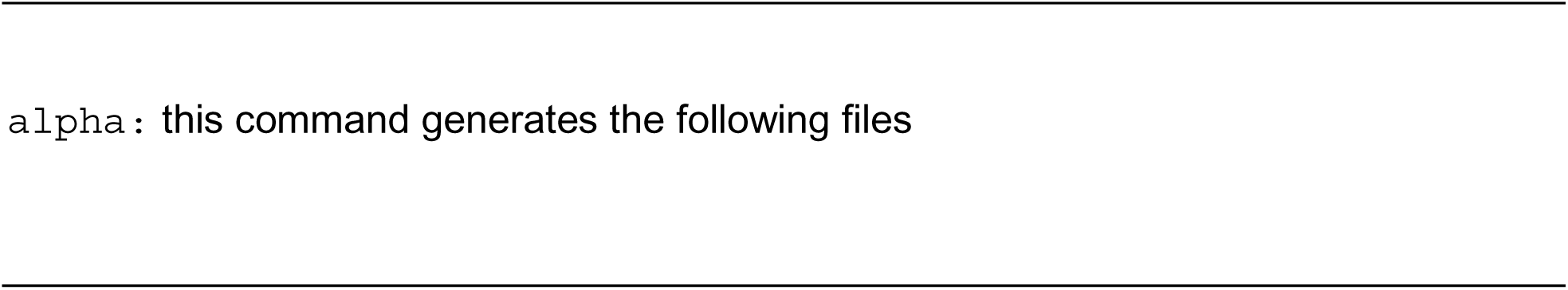

- *alpha_div/calc_[mysample]–[alpha].txt*

- The raw calculations of alpha diversity per sample
- *alpha_div/stats_[mysample]–[alpha].txt*

- Kruskal-Wallis test as a omnibus test, with post-hoc pairwise group tests using Wilcoxon rank-sum test with continuity correction with FDR
- *Plots/alpha_div_[mysample]/[group]–[alpha].svg*

- Violin plots with only statistically significant comparisons indicated (Wilcoxon rank-sum test).

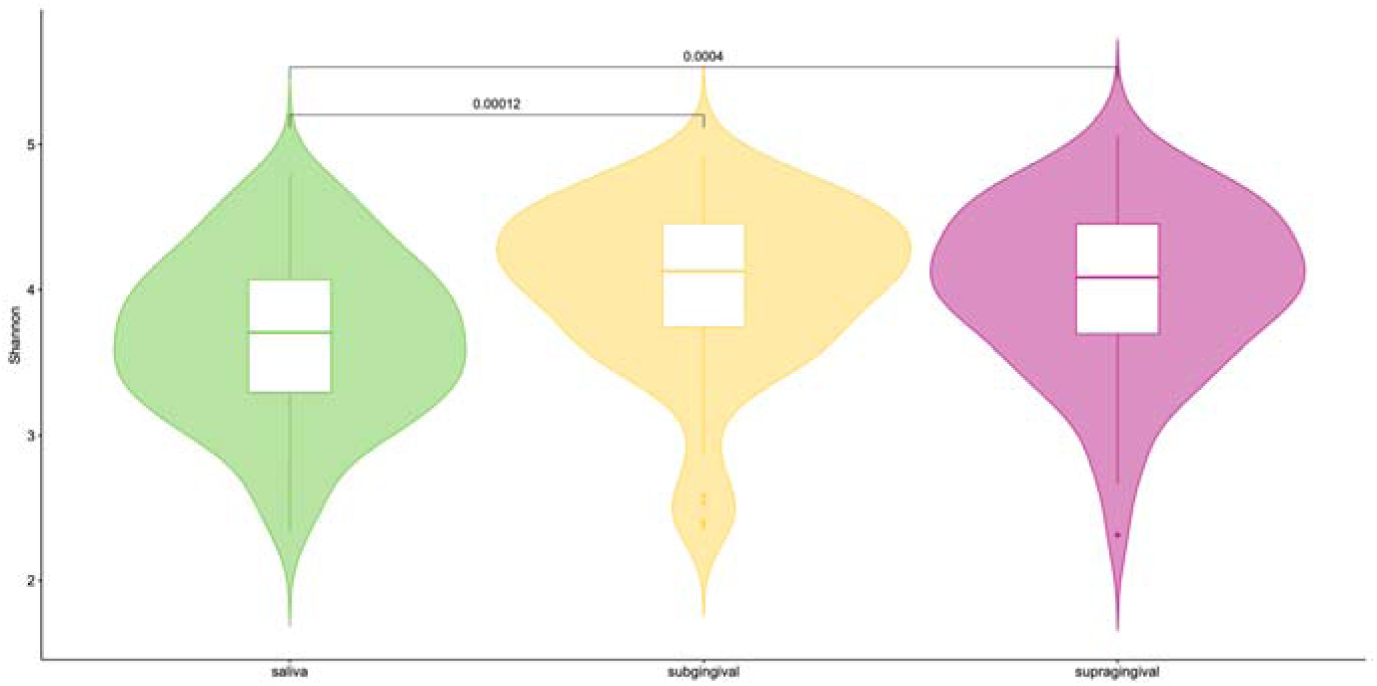

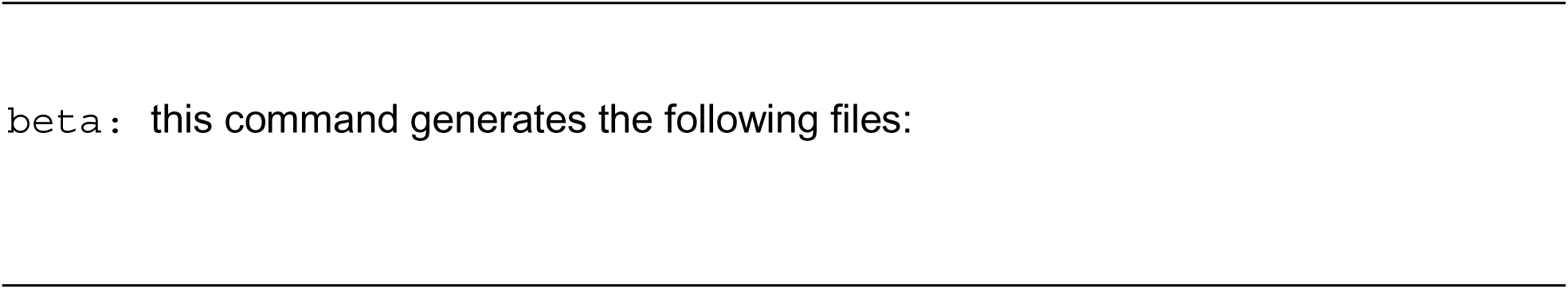

**The raw calculations of beta diversity dissimilarity**

- *distance/beta_div/ [mysample]–[distances].txt*

- The raw calculations of beta diversity dissimilarity

**ANOSIM (ANalysis of SIMilarity) outputs**

- *distance/ANOSIM/[mysample]/anosim–[distances]–[group].txt*

- Omnibus test
- *distance/ANOSIM/[mysample]/anosim–[distances]–[group]_pairwise.txt*

- ANOSIM as a post-hoc test for pairwise comparison with FDR-adjusted p-values

**ADONIS (Permutational Multivariate Analysis of Variance using distance matrices using adonis2 vegan implementation) outputs**

- *distance/ADONIS/[mysample]/adonis–[distances]–[group].txt*

- Omnibus test
- *distance/ADONIS/[mysample]/anosim–[distances]–[group]_pairwise.txt*

- ADONIS as a post-hoc test for pairwise comparison with FDR-adjusted p-values

**Beta dispersion (Multivariate homogeneity of groups dispersions (variances))**

- *distance/PERMDISP/[mysample]/betadisper–[distances]–[group].txt*

- Both the omnibus-test, and the pairwise analysis

**Dunn’s test for multiple comparisons**

- *distance/PerGroupDistCompare/[mysample]/Dunns–[distances]–[group].txt*

- Comparison of the distances per group-pairs.
- Used to test if the distances between three groups are statistically different from each other or not.

- Eg: saliva-subgingival plaque compared to saliva-supragingival plaque tests if the distance between saliva and the subgingival plaque is statistically different from that from saliva to supragingival plaque

#### Plots

**Beta dispersion through Principal Coordinates Analysis (PCoA)**

- *plots/PCoA_betadispersion–[mysample]–[distances]–[group].svg*

- The ellipsoids are drawn with 1 standard deviation of the dispersion, as per Vegan’s default. 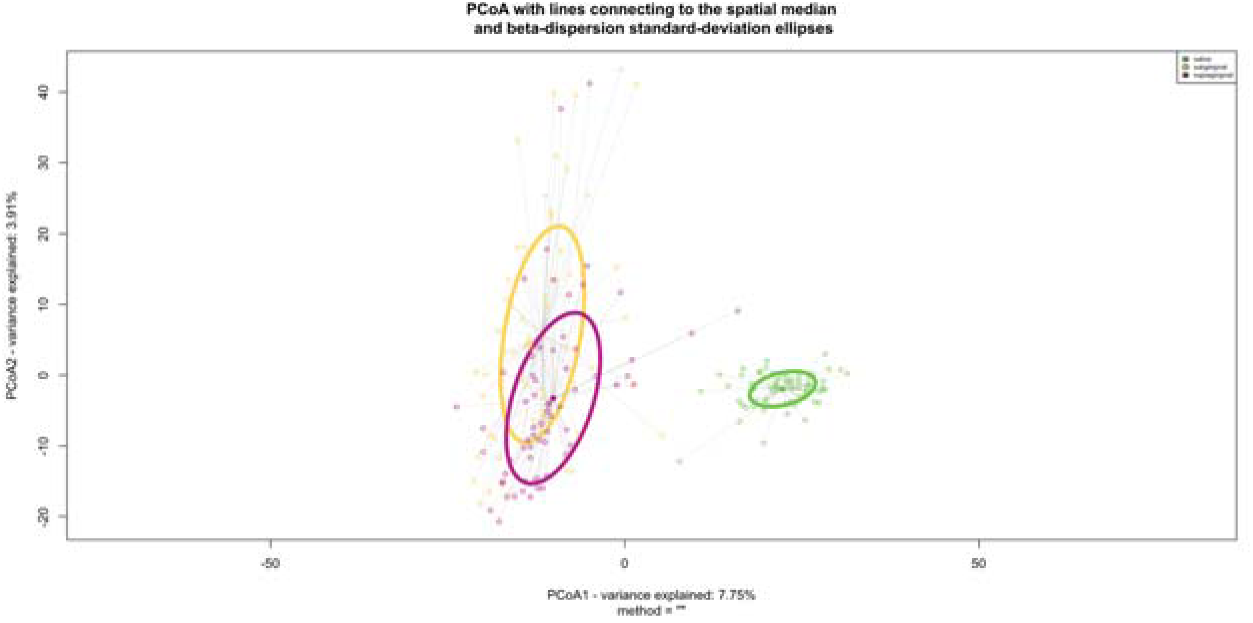
- *plots/boxplot_betadispersion–[mysample]–[distances]–[group].svg*

- Box-plots of the beta dispersion away from the centroids. 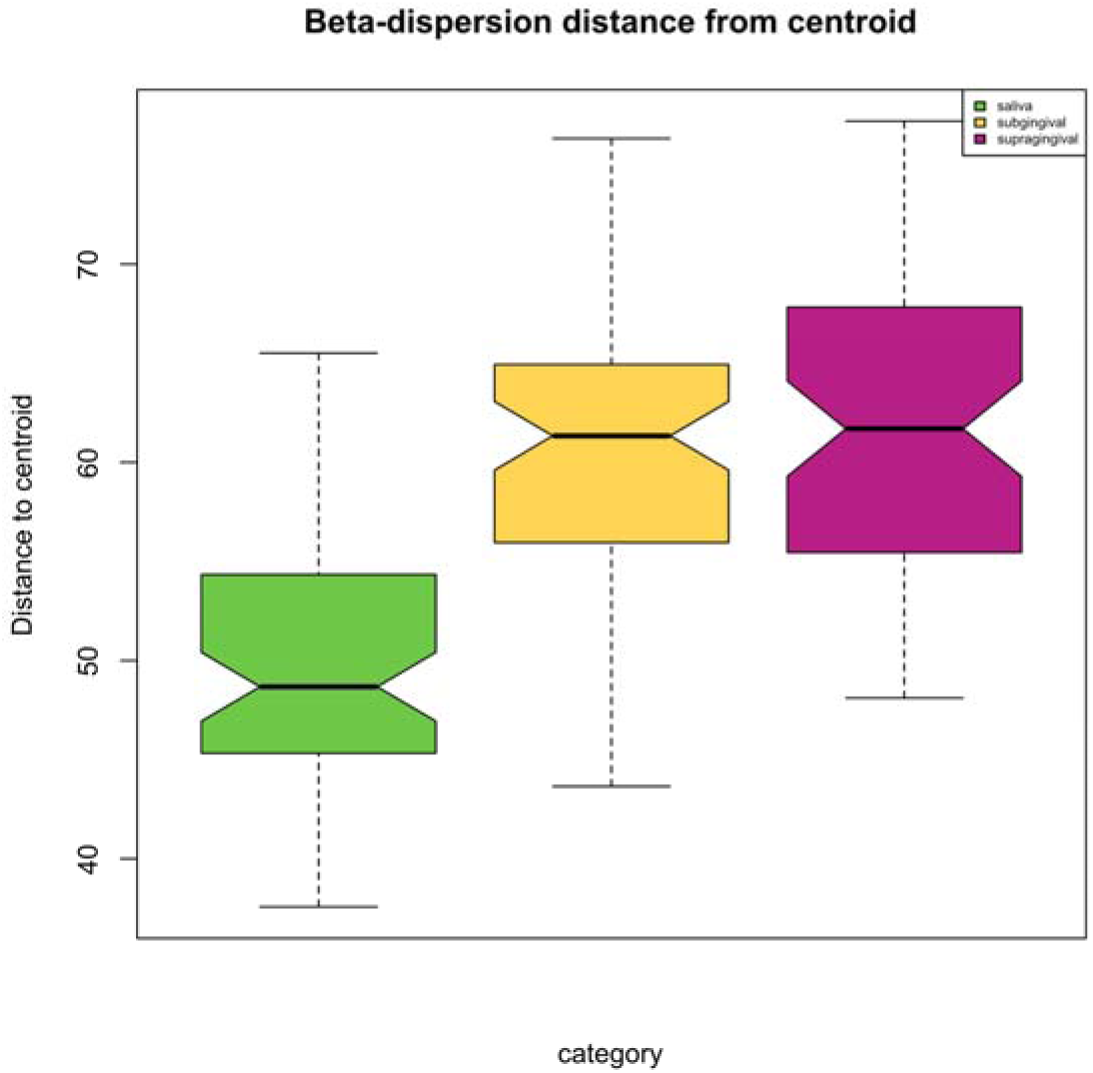

**Principal Coordinates Analysis (PCoA)**

In addition to the PCoA with the beta dispersion above, four different versions of PCoA plots are also generated, depending on the presence/absence of two decorations; probability density plots, and sample names. Plots are as below:

- plots/betaDiv_[mysample]/PCoA–[distances]–[group].svg 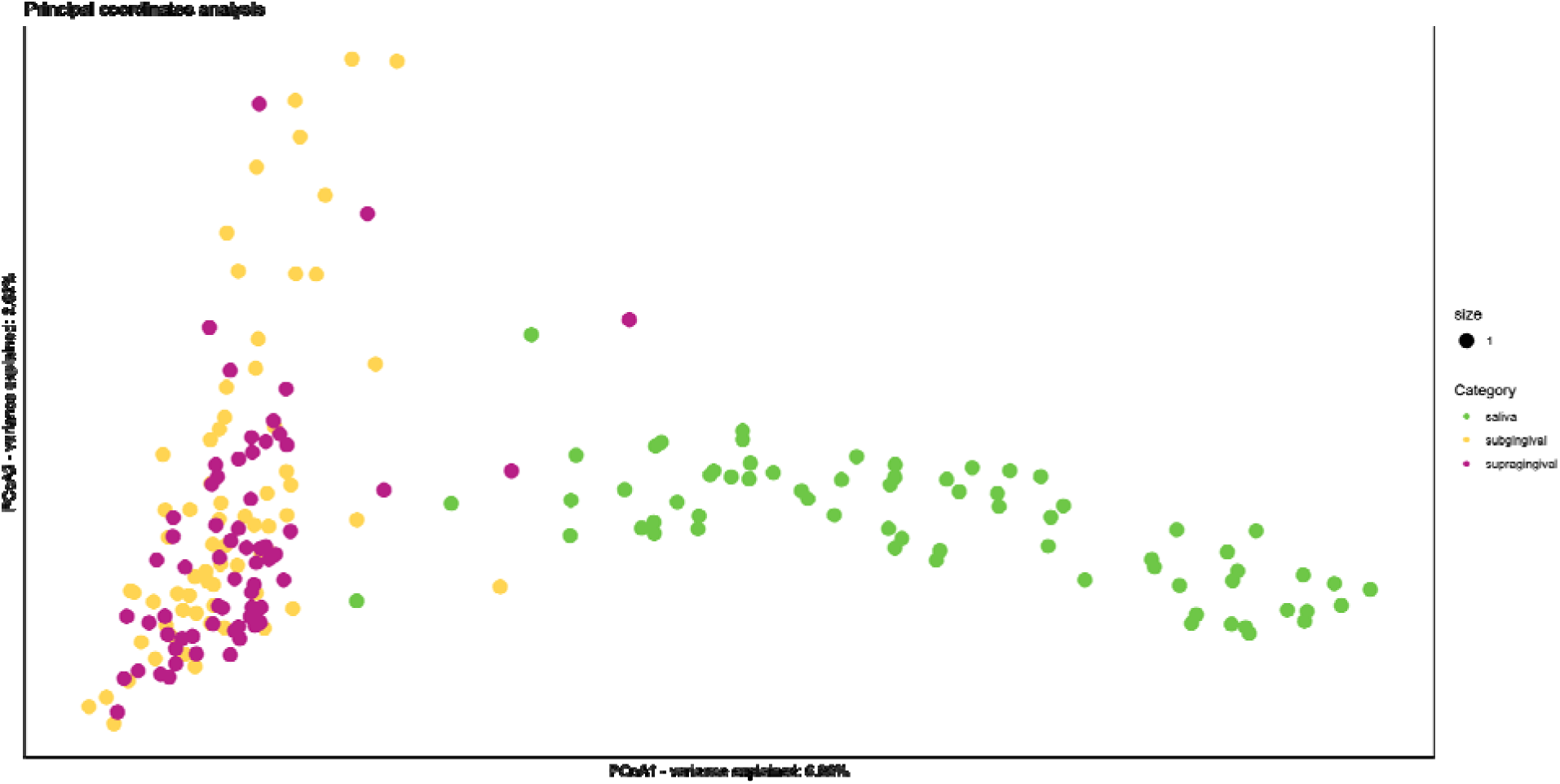
- *plots/betaDiv_[mysample]/PCoA–[distances]–[group].svgnoNameswProbDF.svg*

- Same as the above plot but with probability density functions on the PC1 and PC2 plots. These can be useful when you have 4+ groups being plotted with overlapping dispersion that understanding the distribution of the groups, as seen in the example below 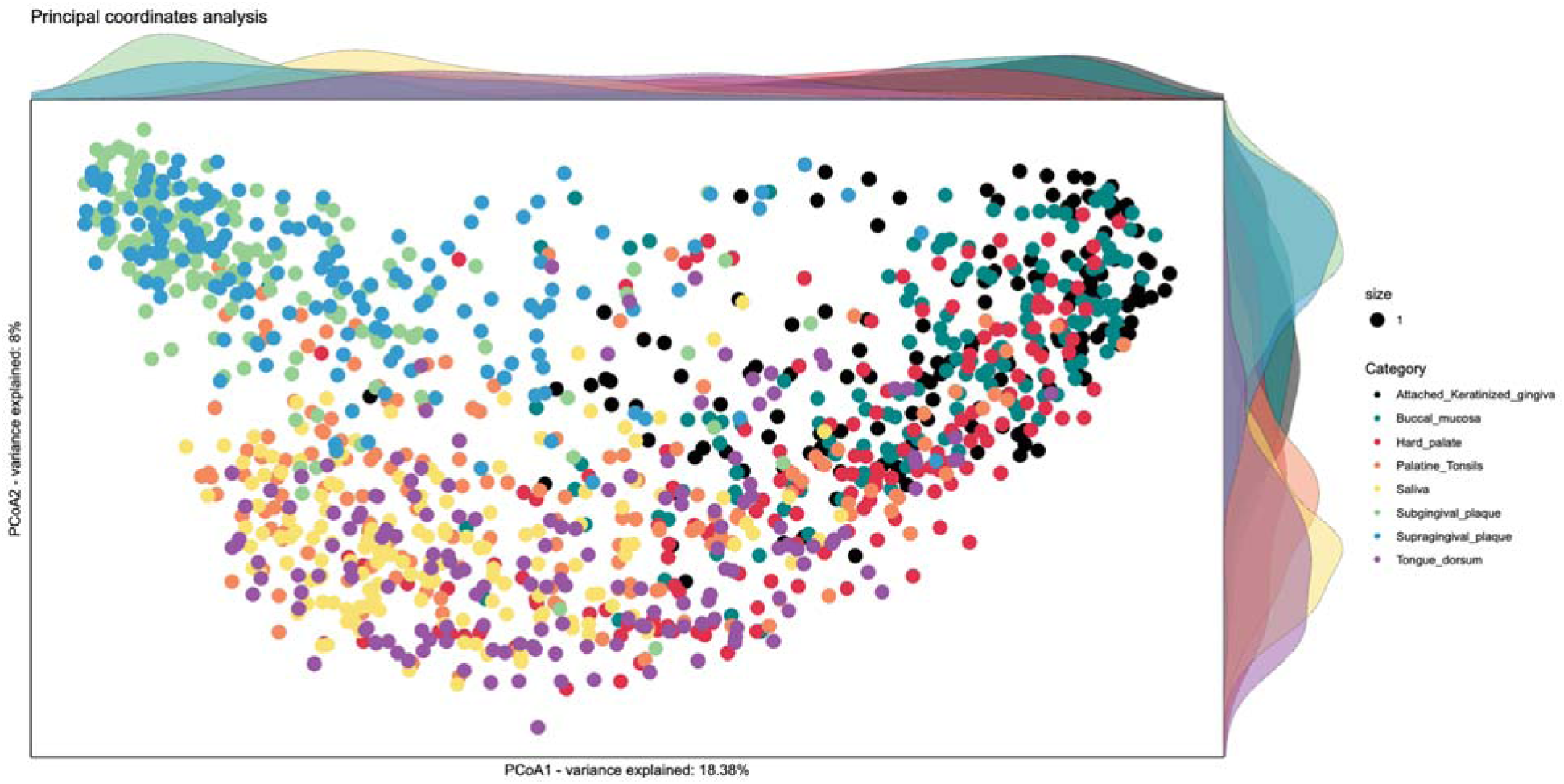
- *plots/betaDiv_[mysample]/PCoA–[distances]–[group].svgwithnames.svg*

- PCoA graph, with sample names. Some overlapping of names is allowed as you can see below. 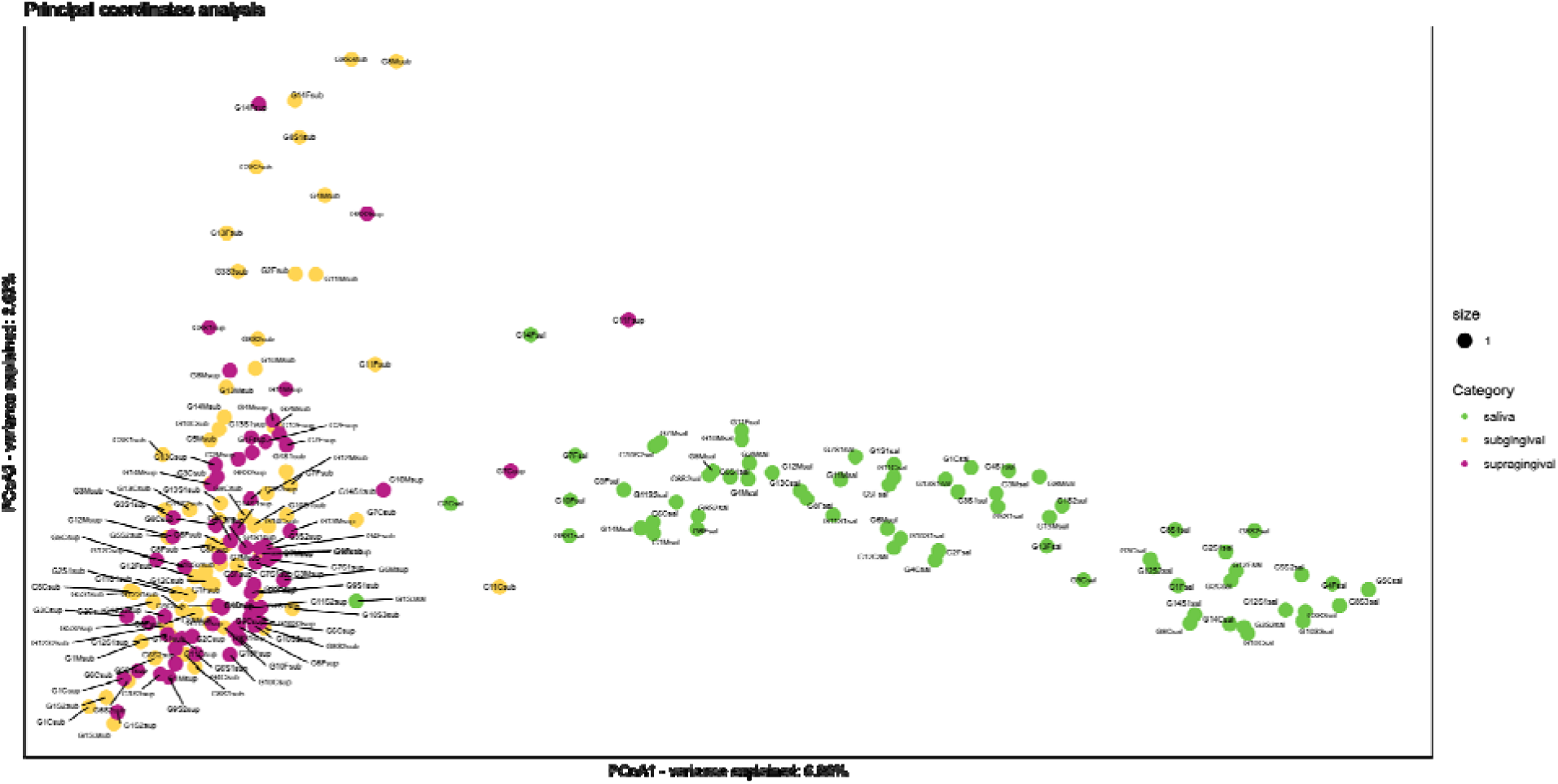
- *plots/betaDiv_[mysample]/PCoA–[distances]–[group].svgwithnamesprobDF.svg*

- Combination of the two decorations above (PDF and sample names) 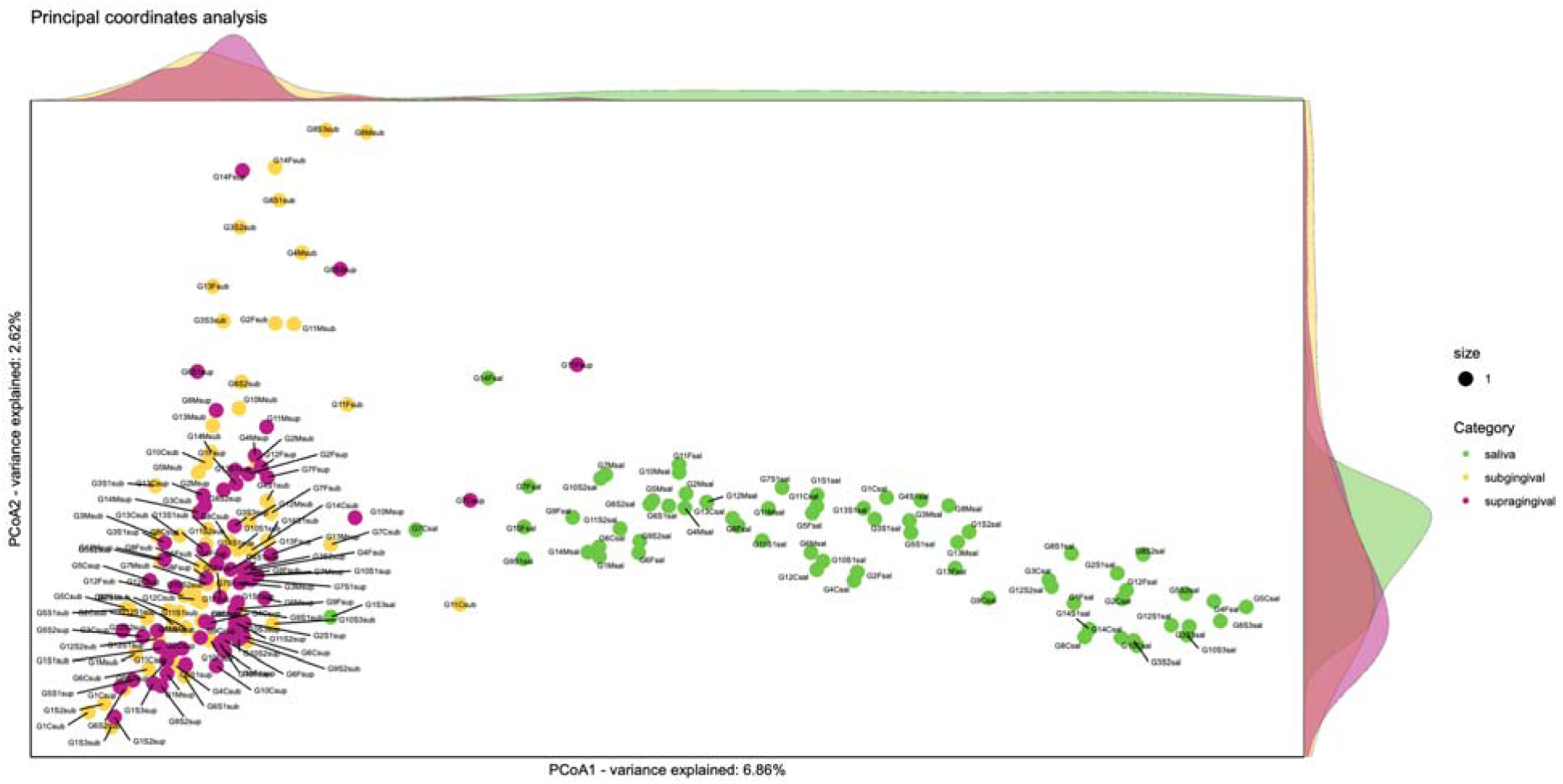

**Non-metric Dimensionality Scaling (NMDS)**

The four plots above are also available as NMDS plots.

- *plots/betaDiv_[mysample]/NMDS–[distances]–[group].svg* 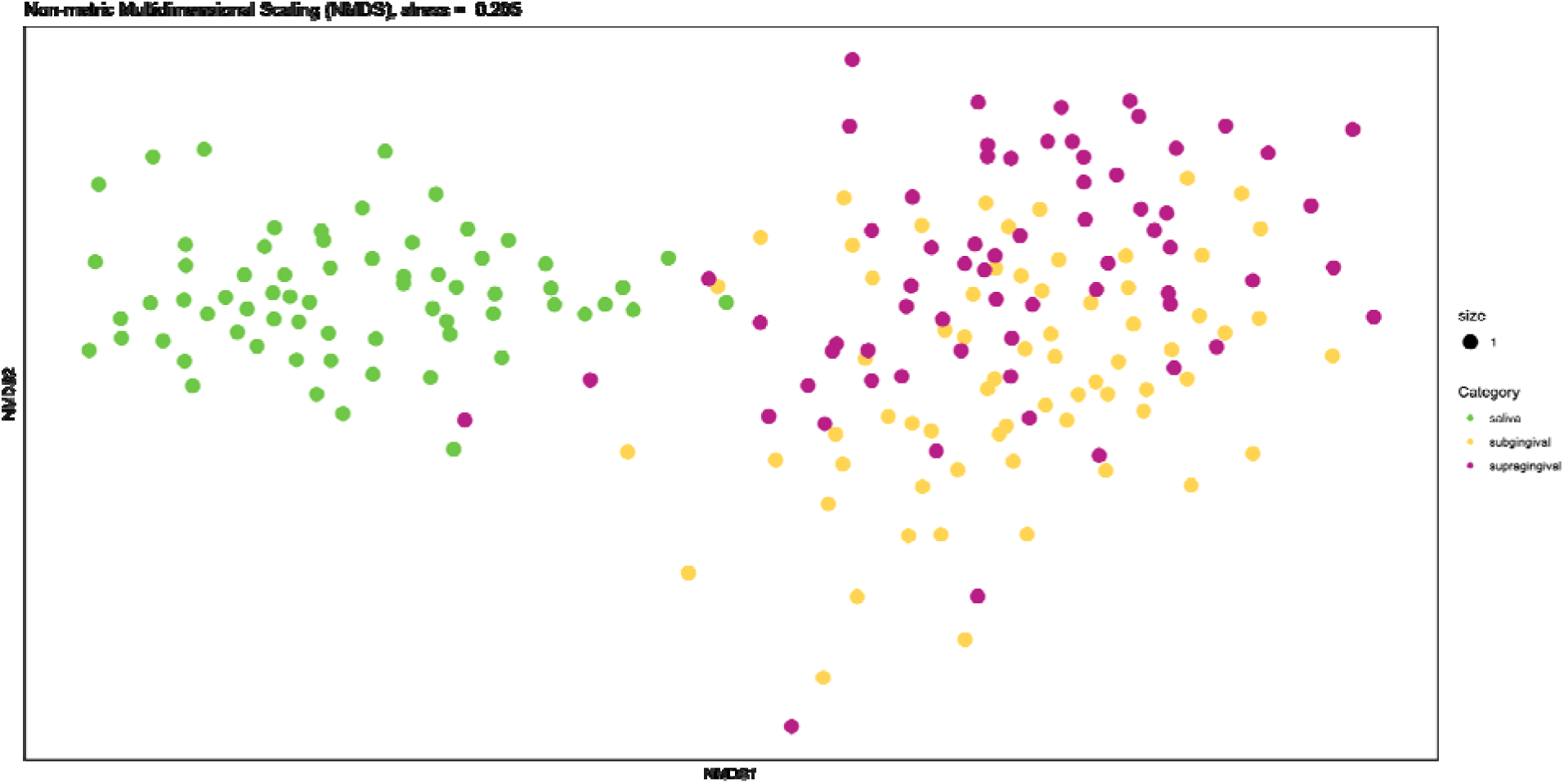
- *plots/betaDiv_[mysample]/NMDS–[distances]–[group].svgnoNameswProbDF.svg* 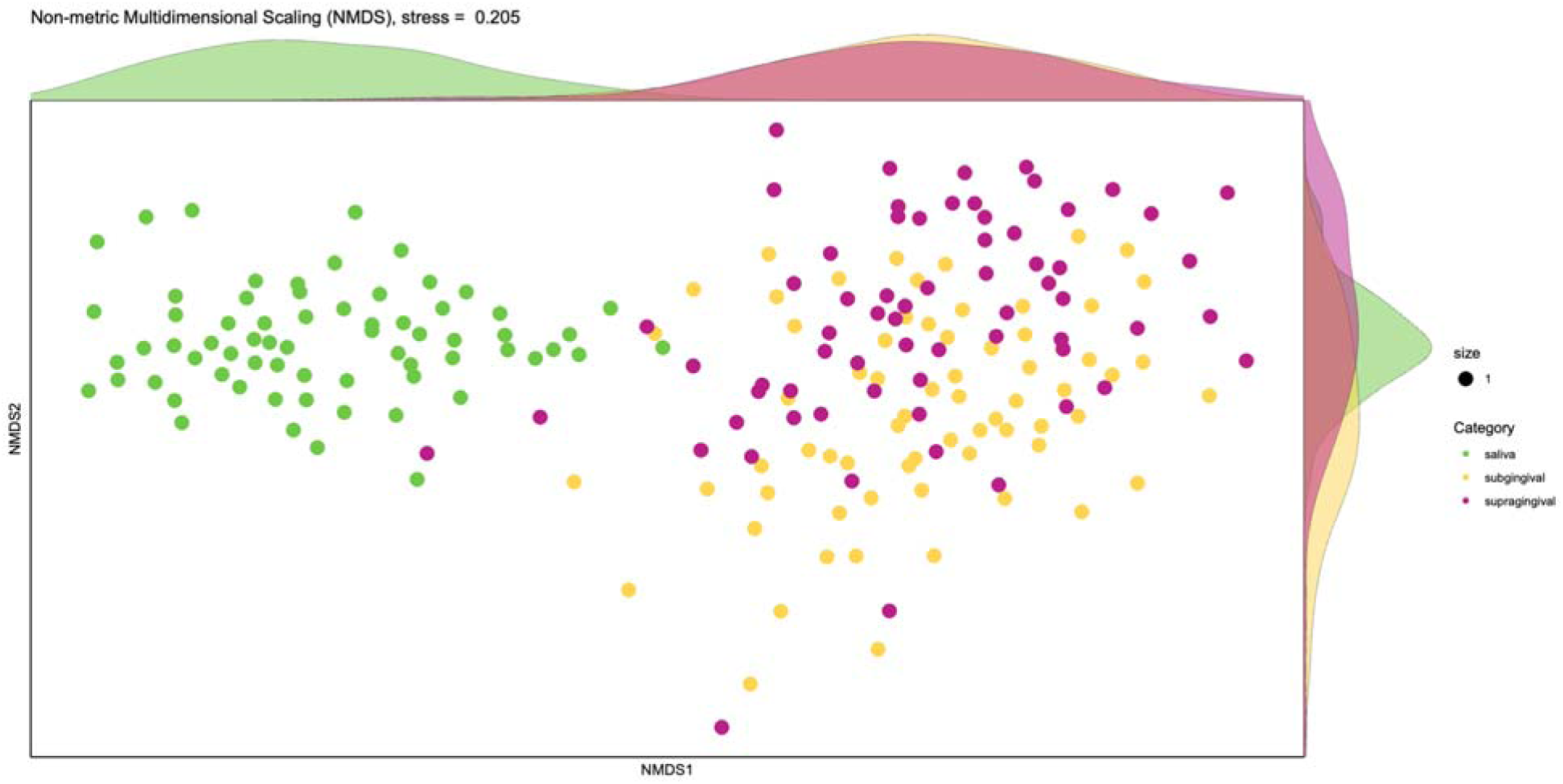
- *plots/betaDiv_[mysample]/NMDS–[distances]–[group].svgwithnames.svg* 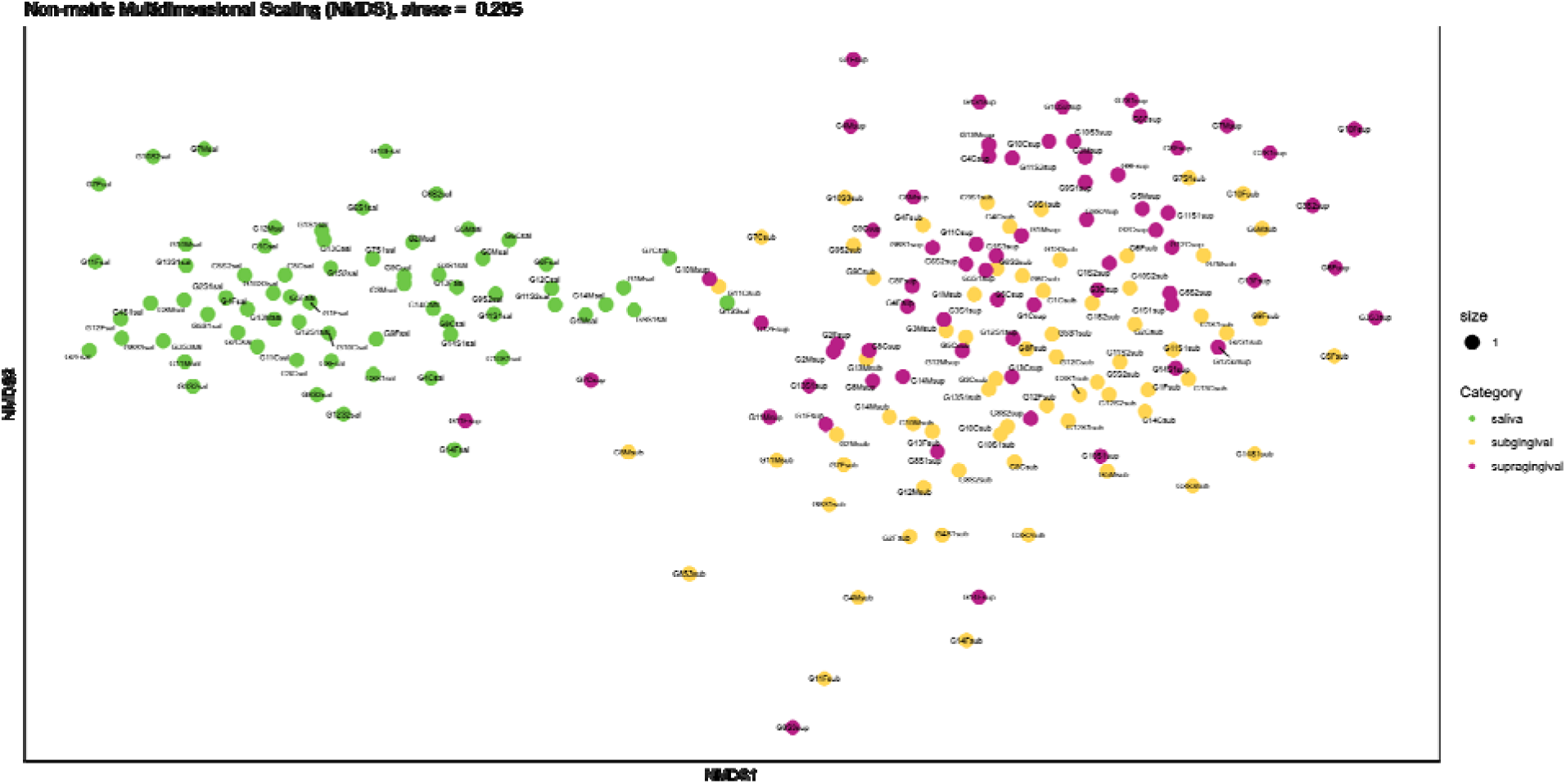
- *plots/betaDiv_[mysample]/NMDS–[distances]–[group].svgwithnamesprobDF.svg* 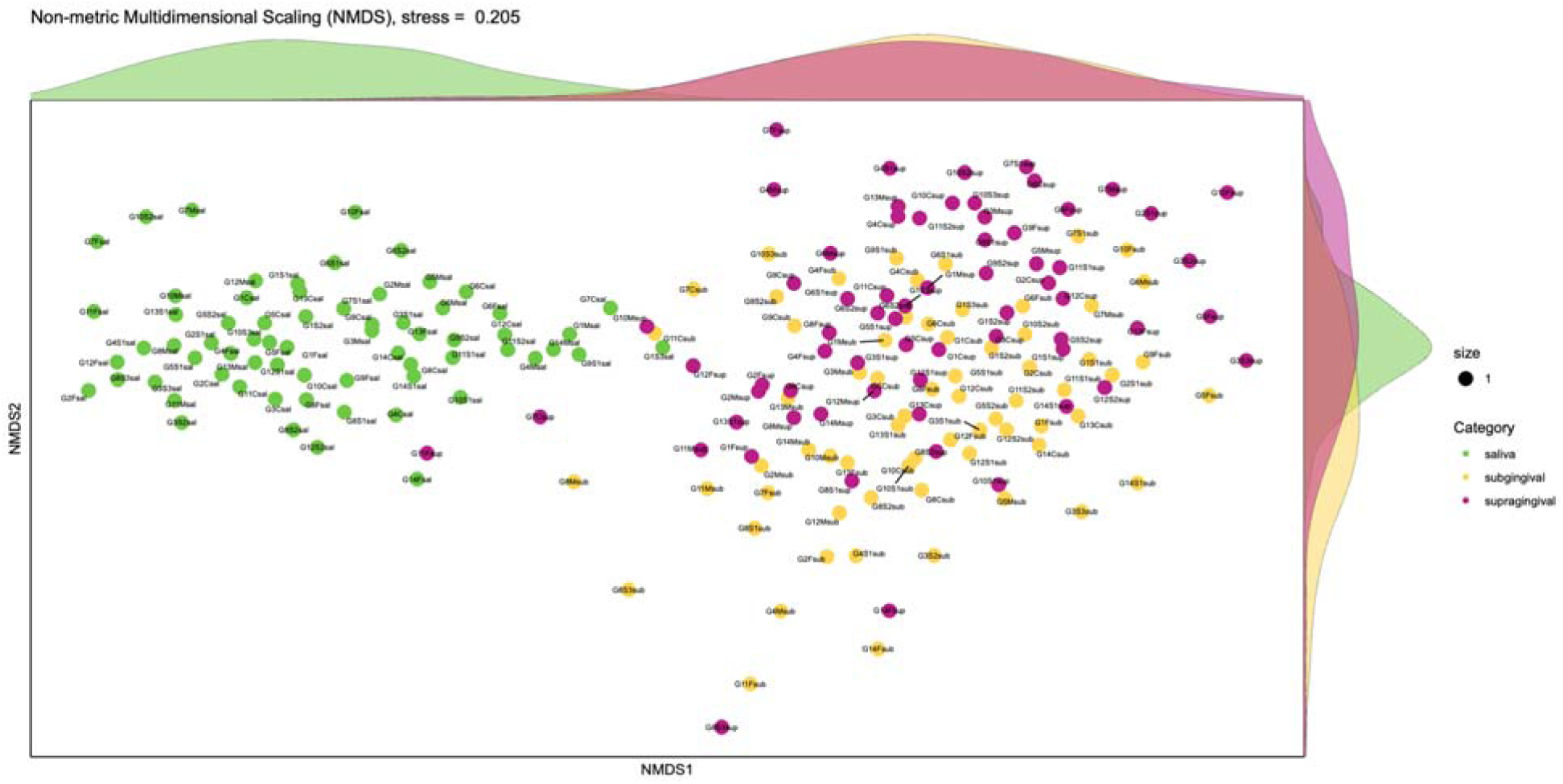

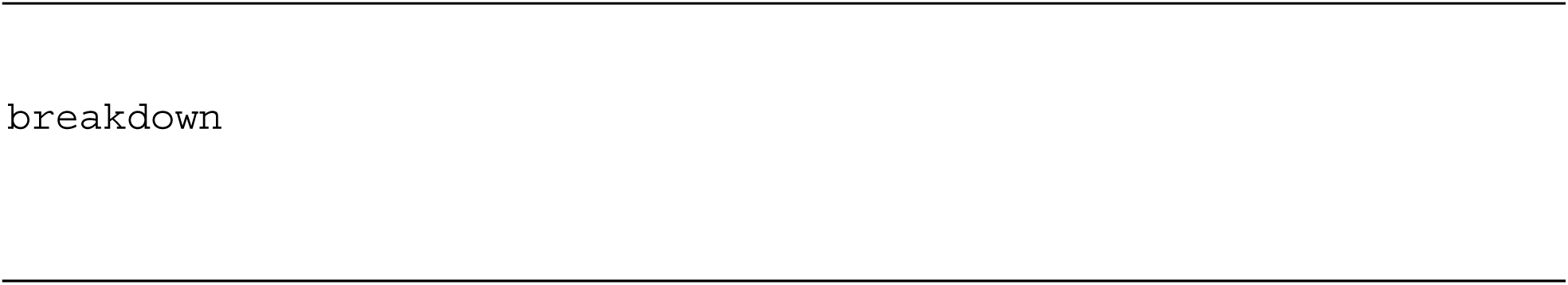

This commands is based on Baselga’s work on breaking down two popular dissimilarity matrices; Bray-Curtis, and Jaccard dissimilarities, between two sites into two components; features that are replaced and those that are not replaced resulting in one site being a superset of another. The `breakdown` command analyzes pairwise samples into these two components. In this command, [distances] is either bray for Bray-Curtis dissimilarity, or jaccard for Jaccard dissimilarity. Generated files are as follows:

- *betapart/[mysample]–[distances]– [category]/ingroupdiff/[category]+[group:group1]-ingroupdiff.txt*

- Results of within-group differences between samples.
- Each category in [group] is generated separately in this folder
- Within-group differences are reported as

- Dissimilarity: raw values, and percentages of the total dissimilarity
- Permutation based mean values: both means, and standard deviations of the distribution are calculated for dissimilarity, and its two components. These are based on the number of permutations [betapart_permutations] and the number of samples to include per permutation [betapart_samples]
- Note: permutation results are saved in *betapart/[mysample]– [distances]–[category]/permuations*
- *plots/Breakdown–[mysample]–[distances]–[category].svg*

- Plot of dissimilarity components of the permutational values 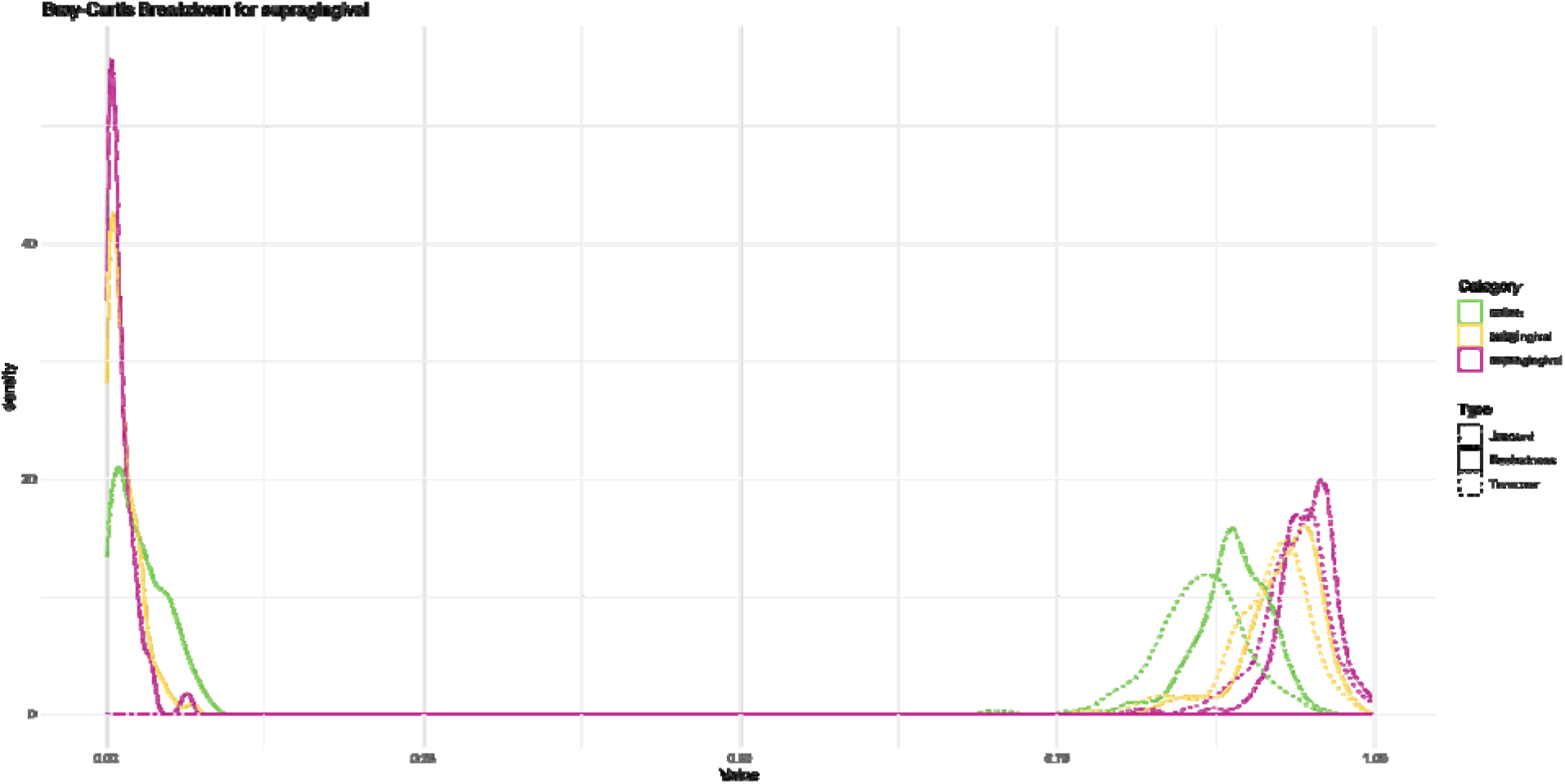
- *plots/betapart_pairwise/[mysample]–[distances]– [category]/[category:group1]+[category:group2].svg*

- An alternate plot for pairwise group comparisons, based on the raw values of the two components.

- Note: a .tsv file is saved in the same folder for the raw values 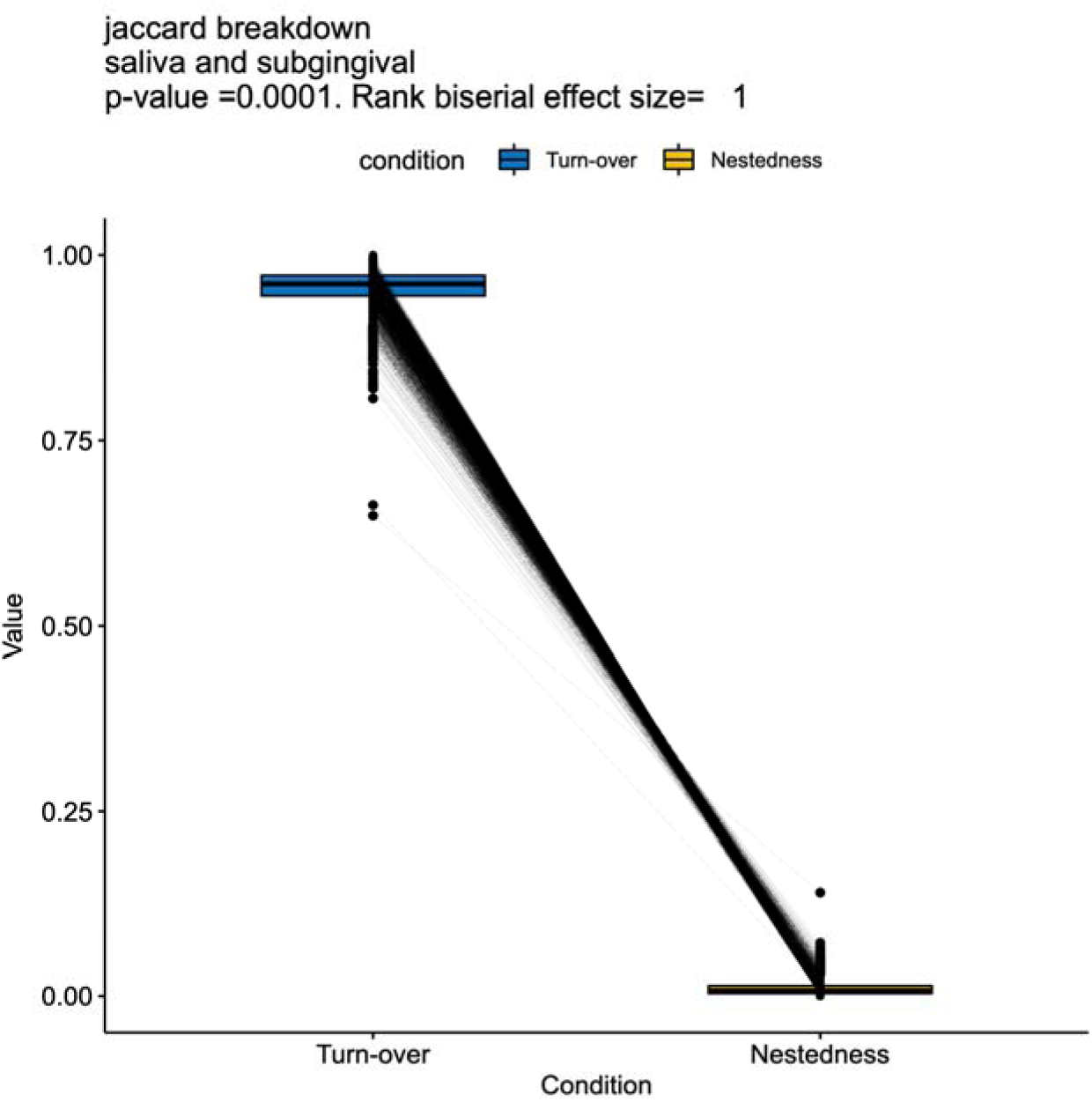

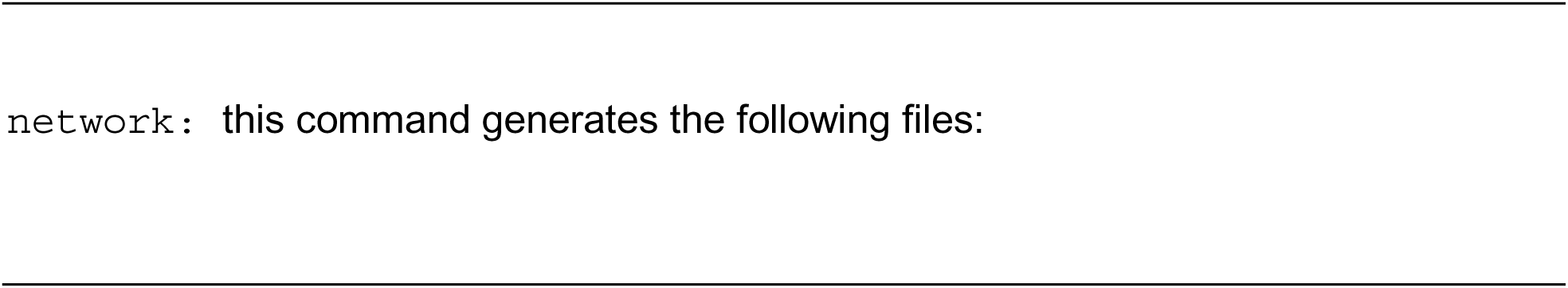

This command generates network plots based on SparCC. The following will explain the steps done to generate the results, and what parameters in the falaphyl.yaml file are used.

**<u>Step 1: Core filtering</u>**

Filtering biom files to features available in each group in [category] to only those features that are present in [threshold] or more %. Core files are saved in:

- *network/[mysample]–[category]/core/[category:group1].tsv*

**<u>Step 2: SparCC calculations</u>**

Calculations are done for all pairwise samples. Files are saved in:

- *network/[mysample]–[category]/corr/[category:group1].tsv*

P-value calculations are done based on the specified number of permutations [sparcc_bootstrap]. The calculations are saved in:

- *network/[mysample]–[category]–corr/pvalue–[category:group1].tsv*

**<u>Step 3: Filtering and ZiPi value calculations</u>**

Filtering is done based on the desired level of relationship strength [sparcc_corr], and p-value [sparcc_pvalue].

Next, modularity is calculated using Louvain clustering from the R package igraph. These are used to calculate the importance of the nodes through its within module connectivity (Zi) and among modules connectivity (Pi). The saved files are formatted to be readily imported into Gephi as they are saved as nodes and edges files in the below locations.

- *network/[mysample]–[category]/nodes–[category:group1].tsv*
- *network/[mysample]–[category]/edges–[category:group1].tsv*

**<u>Step 4: ZiPi plotting</u>**

ZiPi values are plotted. Files are found in the following locations:

- *plots/network/ZiPi_[mysample]–[category]/ZiPi–[category:group1]*

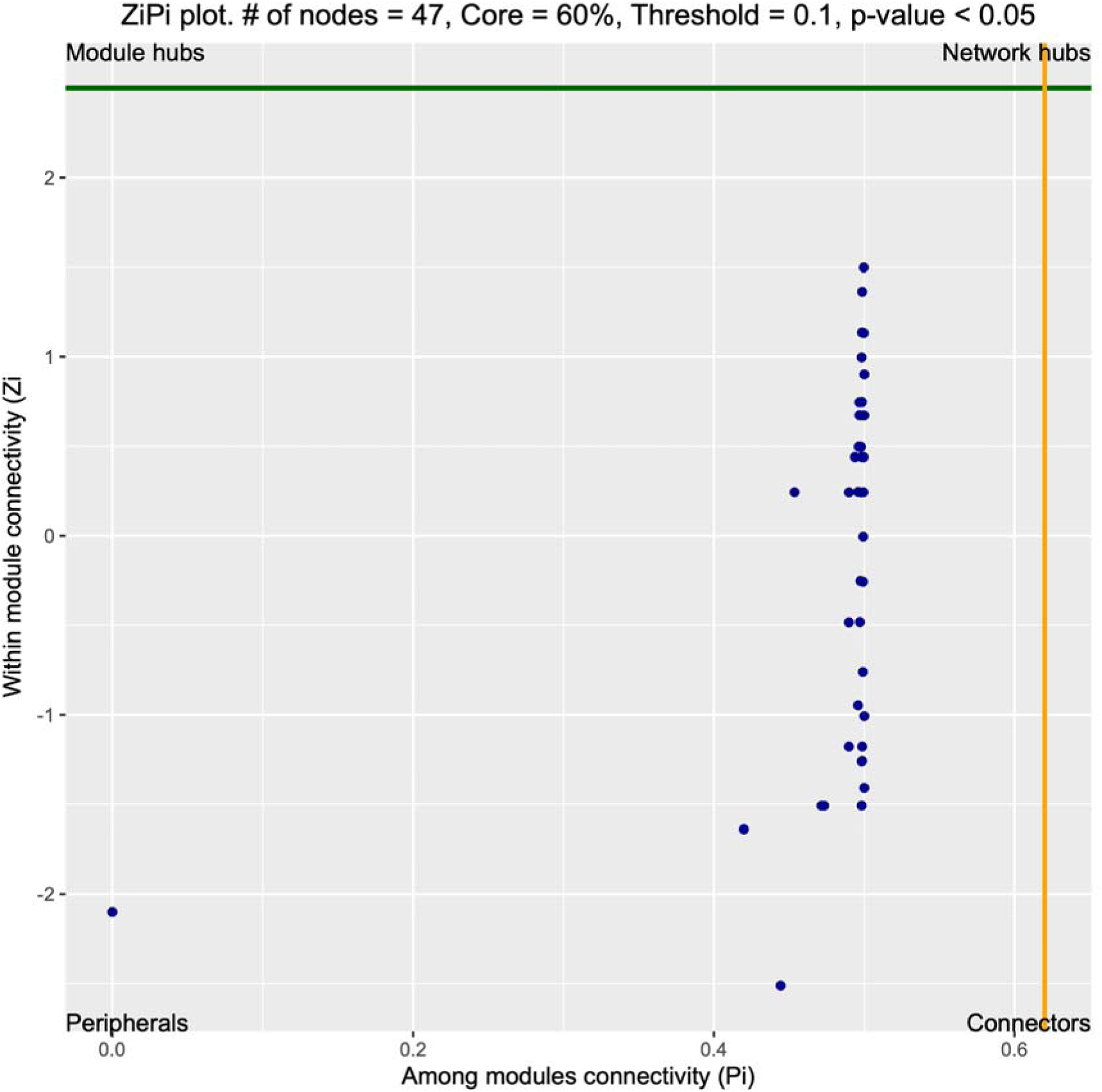

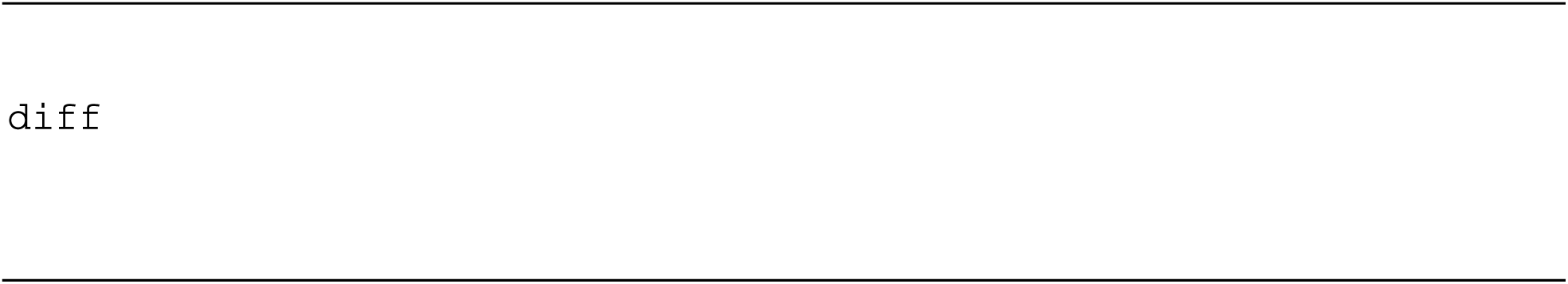

This command generates differential abundances between groups in [category]. The command heavily uses the R package DAtest with some minor in-house changes that are explained below. To explore all the different differential abundance features that are available in this pipeline, refer to the falaphyl.yaml file under the [DA_tests] section. DA calculation is performed as follows:

**<u>Step 1: Feature prefiltering</u>**

Features are prefiltered prior to differential abundance calculations. Features that do not meet the following criteria are grouped together as a feature named “Other”. This is done to prevent loss of data that may skew the results due to the compositionality of the microbiome data.

- [DA_minsample]: The minimal number of samples a feature must be present for it to be retained.
- [DA_minread]: The minimal number of reads a feature must be present for it to be retained.
- [DA_minabund]: The fraction of the minimal mean relative abundance a feature must occupy for it to be retained.

**<u>Step 2: Differential abundance calculations</u>**

Each one of the tests in [DA_tests] is calculated separately after filtering. If [category] contains 3+ groups, then calculations are automatically generated for each pairwise groups.

In addition to the methods listed in [DA_tests] that are natively supported within the package, FALAPhyl also supports Compound Poisson Linear Models (CPLM) through its implementation in Tweedieverse. Calculations are saved in the below location:

- *diff/[mysamples]–[category]– minAbd[DA_minabund]minR[DA_minread]minS[DA_minsample]/diff– [category:group1]![category:group2]–[da_tests:test1].tsv*

**<u>Step 3: Area Under the Curve, False-Discovery rate, and Power calculations</u>**

DAtest package automatically calculates the suitability of each test for the dataset through its resilience in detecting the differential abundances if the data were spiked. FALAPhyl parallelizes the step to generate each spike, defined under [DA_effectsize]. Each spike is repeated 20 times.

Spike-in test results are saved in the following files:

- *diff/[mysamples]–[category]– minAbd[DA_minabund]minR[DA_minread]minS[DA_minsample]/AUC_FDR_Power/effe ctSize–[DA_effectSize]–[DA-tests].tsv*

**<u>Step 4: Plotting the graphs</u>**

The original plots of DAtest have been reimplemented. The following are the generated graphs with what was changed from the original DAtest implementation:

- *plots/Score–[mysamples]–[category]– minAbd[DA_minabund]minR[DA_minread]minS[DA_minsample].svg*

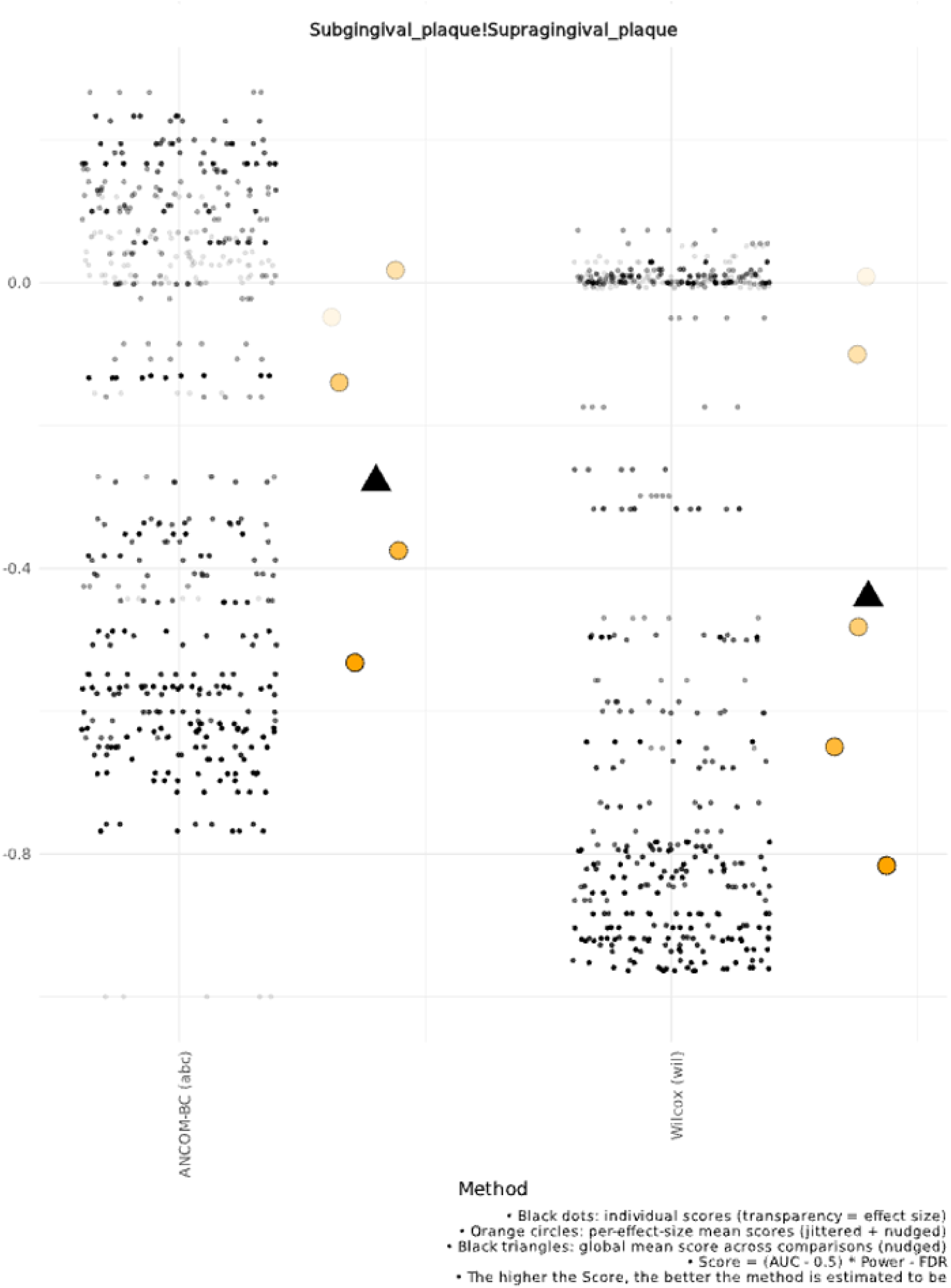

The averages of each spike used for the test’s effect size strength is added in the plot.

Moreover, average of all spike tests is also added. Explanations of how scores are calculated are added to the bottom of the plot.

- *plots/Power–[mysamples]–[category]– minAbd[DA_minabund]minR[DA_minread]minS[DA_minsample].svg*

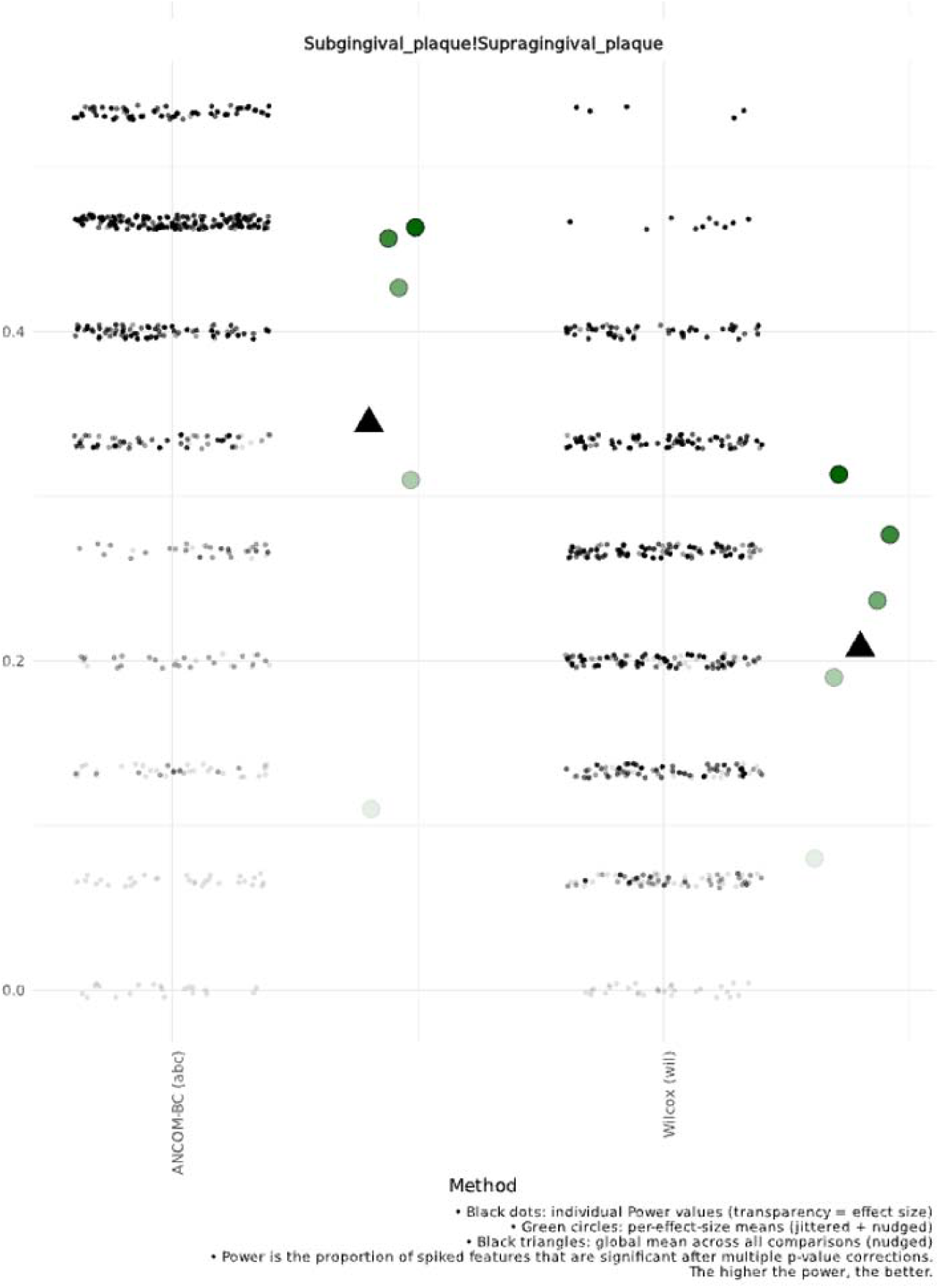

The averages of each spike used for the test’s effect size strength is added in the plot. Moreover, average of all spike tests is also added. Explanations of what power means in terms of a spike test is added.

- *plots/FDR–[mysamples]–[category]– minAbd[DA_minabund]minR[DA_minread]minS[DA_minsample].svg*

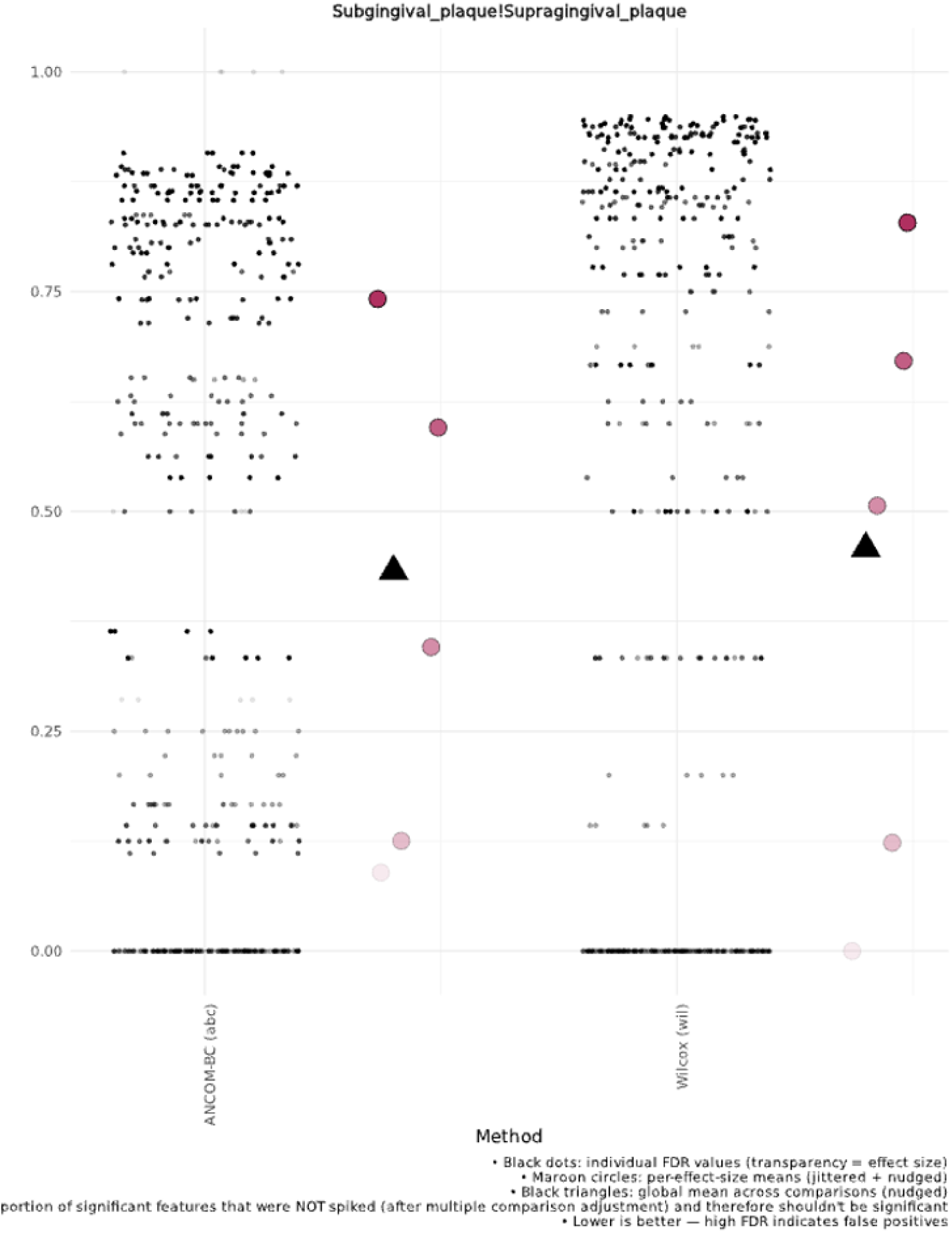

The averages of each spike used for the test’s effect size strength is added in the plot. Moreover, average of all spike tests is also added. Explanations of what FDR means in terms of a spike test is added.

- *plots/AUC–[mysamples]–[category]– minAbd[DA_minabund]minR[DA_minread]minS[DA_minsample].svg*

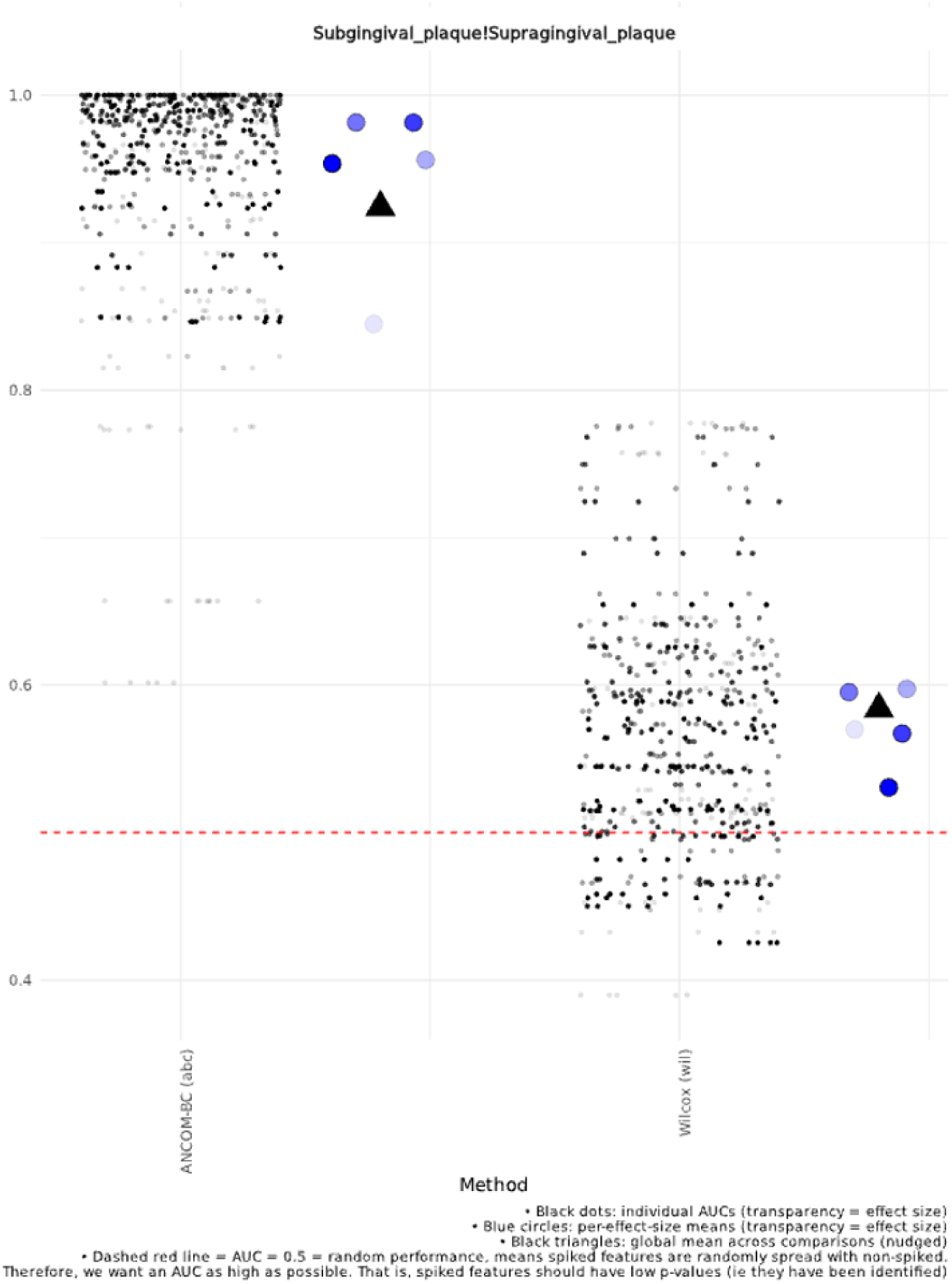

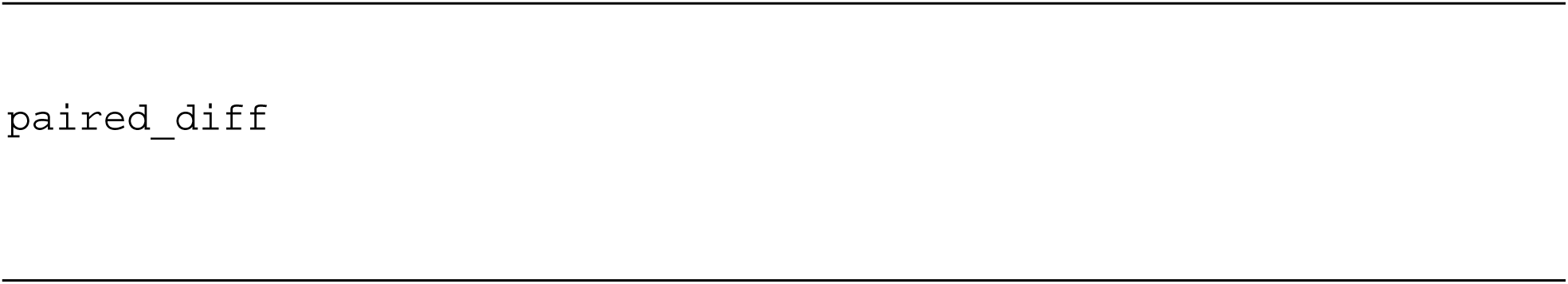

This command generates the same results as the above command, with the difference is that the groups are done at a sample-level, instead of group level. This is done by filtering [category] samples to only those that are present in [subjectID]. The files are saved in the same paths as above, with the only difference in the paths having the word “paired” in them.

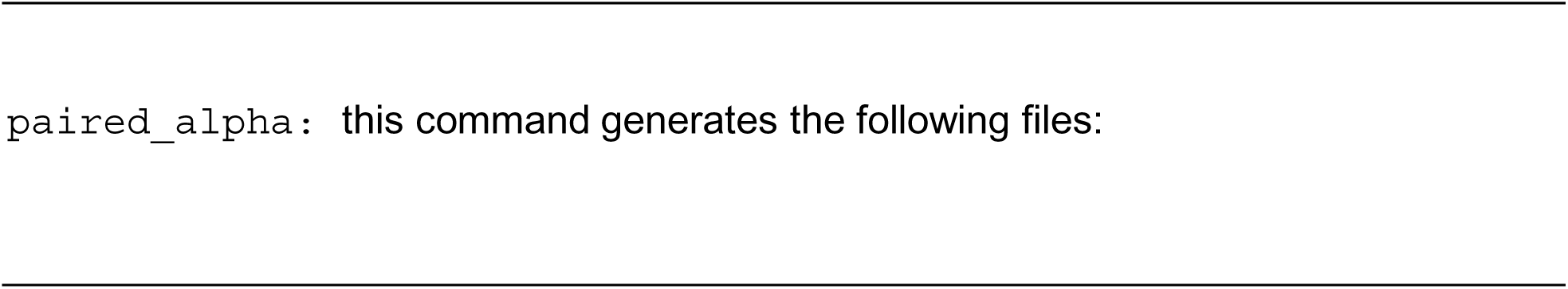

- *plots/patientlvl_alphaDiv_[mysample]–[category]– [alpha]/[category:group1]_[category:group2].svg*

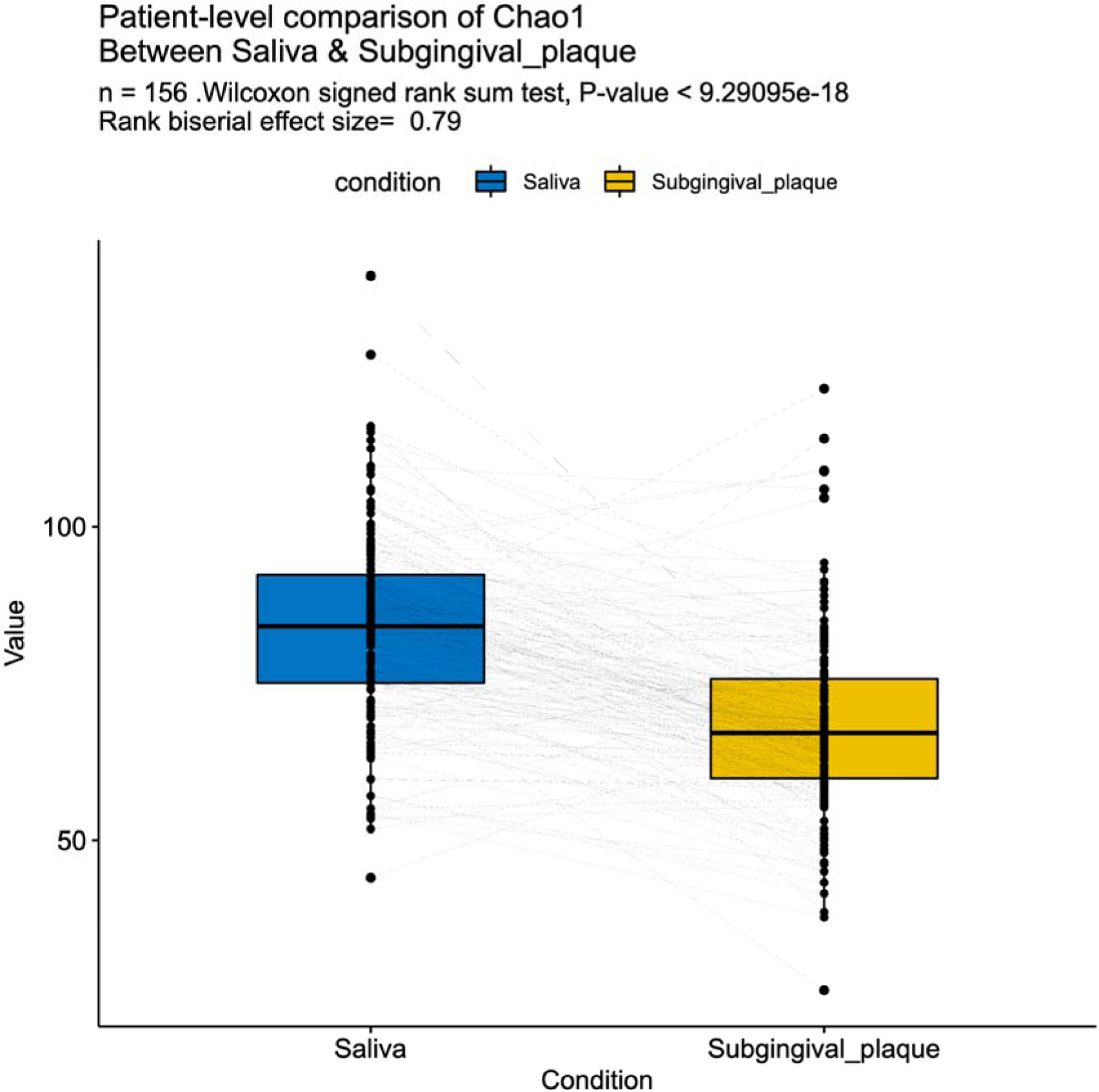

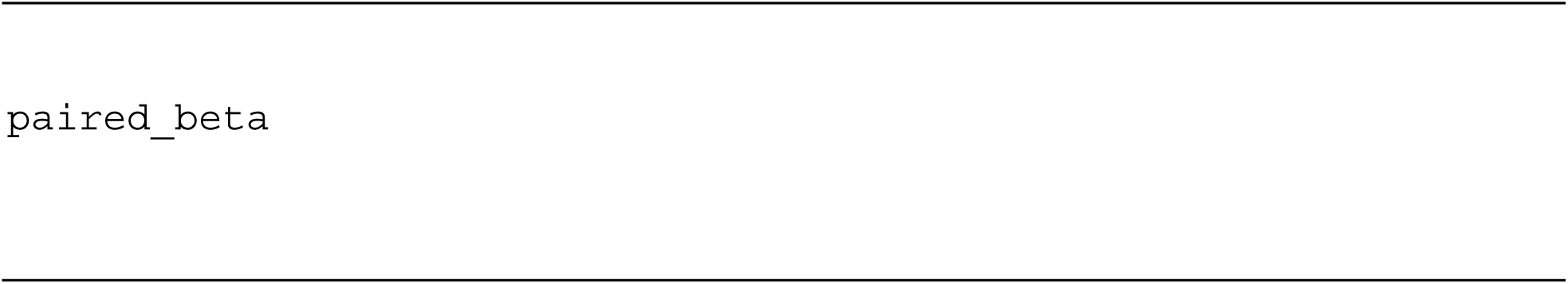

This command does the same analysis as breakdown per group, but restricted to only within the same [subjectID].

- *plots/patientlvl_alphaDiv_[mysample]–[category]– [distances]/[category:group1]_[category:group2].svg*

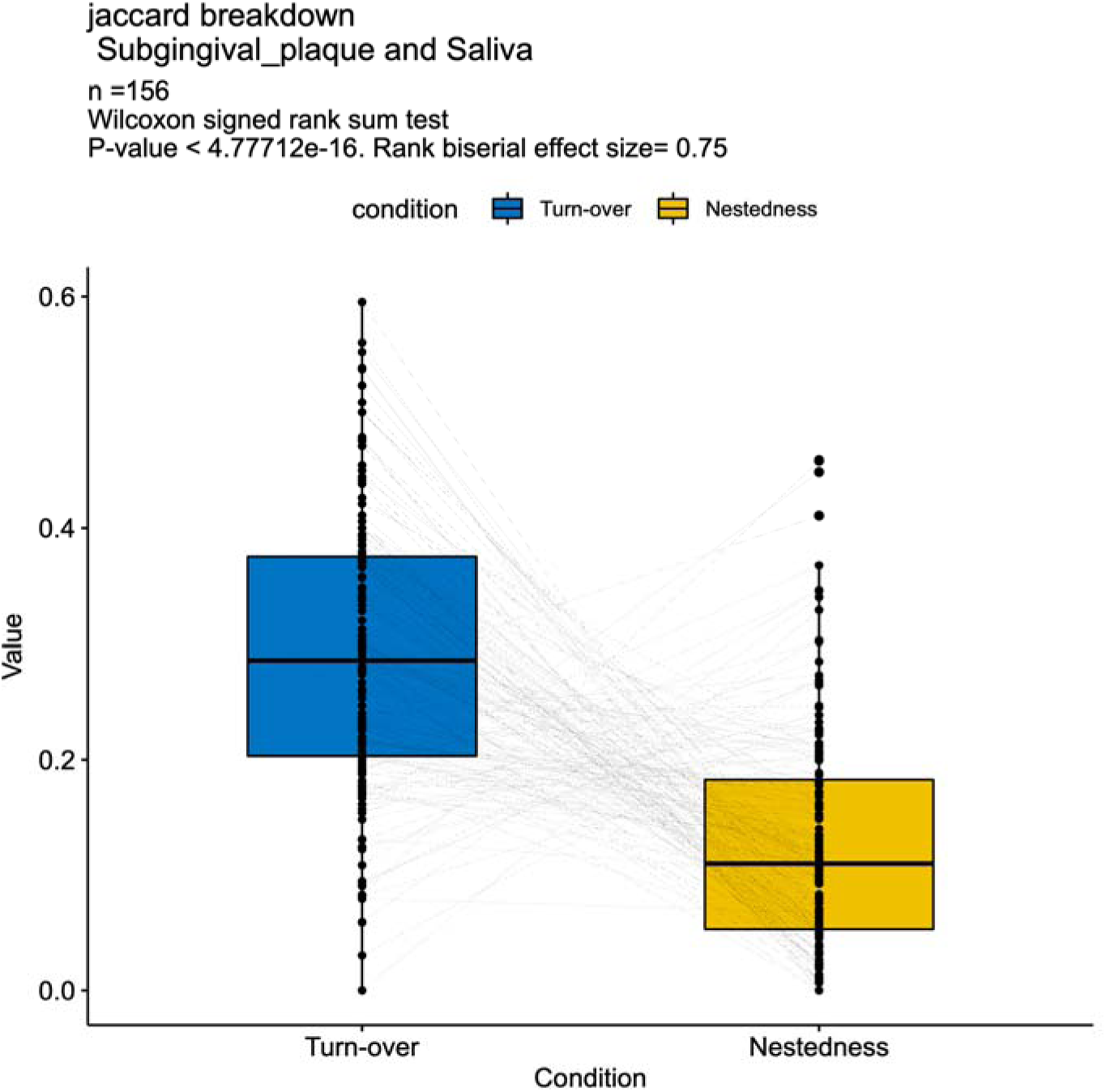

### Appendix 2

#### Case study 2 materials and methods

**Study Design and Participants** The study was approved by the University of Alberta Research Ethics Board (Project No. Pro00120887). This 1-week interventional cohort pilot study recruited 14 families, each containing at least one preschool-aged child (five years old or younger) and at least one sibling. We chose this age for the youngest child to reduce outside-world contact associated with attending schools. Families were recruited from local community organizations such as churches and parent groups.

**Sample Collection** During the initial visit to the Oral Health Clinic at the University of Alberta, samples were collected from various oral sites of all family members. Participants were advised not to eat/drink for at least 30 minutes prior to the appointment. The following procedures were used for sample collection:

- Saliva: Unstimulated saliva samples were collected from adults using sterile tubes. The Micro•SAL™ device (Oasis Diagnostics, Vancouver, Washington, USA) was used for children. The device was placed in the mouth for a set period to absorb saliva when the sampler was saturated, after which saliva was extruded from the cotton and stored at −20°C.
- Buccal Mucosa and tongue: The inside of the cheek (buccal mucosa) was sampled using the DNA•SAL™ device (Oasis Diagnostics). Participants were asked to scrub the buccal mucosa and store it in the solution provided with the device at −20C. Cells were removed from the device by vortexing in 1X PBS and lysozyme for 1 hour. The supernatant was then used for downstream DNA isolation. Tongue samples were collected similarly.
- Supragingival Plaque: Supragingival plaque was collected using sterile paper points, which were gently placed at multiple supragingival sites and held in place for 20 seconds. All collected points were pooled in a single tube containing RNAlater for storage at −20°C.
- Subgingival Plaque: Subgingival plaque was collected from the distal surfaces of the lower central incisors using sterile paper points. The points were inserted subgingivally into the sulcus for 20 seconds, then transferred to a tube containing RNAlater for preservation and storage at −20°C. Following the baseline sample collection, each family’s test subject underwent professional oral prophylaxis to disrupt the existing biofilm in the oral cavity. This procedure involved a comprehensive cleaning of the teeth and gums using rubber cups and pumice to disrupt and remove the existing biofilm, effectively displacing the oral bacteriome. A questionnaire (supplementary) was given to the parents to document familial demographic characteristics, and close contact activities among the family members. Subgingival samples were collected from the test participant one week after the prophylaxis using the same procedure described above.

**DNA Extraction and Sequencing** DNA was extracted from the collected samples using the QIAGEN QIAamp DNA Mini Kit (QIAGEN, Germantown, MD, USA), following the manufacturer’s protocol. The concentration and purity of the extracted DNA were assessed using a Qubit 4 Fluorometer (ThermoFisher Scientific, Waltham, MA, USA) with the Qubit™ 1x dsDNA High-Sensitivity Assay Kit, ensuring that the DNA was of sufficient quality for subsequent sequencing. The V1-V3 regions of the 16S rRNA gene were amplified using specific primers (27F: 5’-ACACTCTTTCCCTACACGACGCTCTTCCGATCTGAAKRGTTYGATYNTGGCTCAG-3’and 519R: 5’-GTGACTGGAGTTCAGACGTGTGCTCTTCCGATCTACGTNTBACCGCDGCTGCTG-3’). The amplified DNA was then sequenced at the Genome Quebec core facility. FASTQ files submitted to NIH SRA (PRJNA1159177).

### Appendix 3

#### Case study 3 materials and methods

##### Participants and Study Design

Participants were recruited from the general community and from the student body of the Faculty of Dentistry at Dentistry Dalhousie University in Halifax, Nova Scotia, Canada to participate in a pilot, parallel arm randomized clinical trial. Study protocol review and ethical approval were obtained from the Dalhousie University Research Ethics Board (REB# 2024-7166). The study protocol and plan for analyses were registered as a clinical trial at ClinicalTrials.gov (identifier: NCT06588049) prior to recruitment and study commencement. Sequences are deposited in NIH SRA (project ID PRJNA1300299).

Recruitment efforts included email invitations and printed posters circulated within the Faculty. Eligible participants were healthy individuals classified as ASA I or II, aged 18 to 40 years on the day of follow-up. Exclusion criteria included pregnancy, uncontrolled systemic diseases or acute infections, use of probiotics, systemic antibiotic or antimicrobial mouthwash use within the past month. Participants with documented allergy to soy, nuts, seeds, dairy, egg products, fish, shellfish or wheat were excluded. The presence of acute or chronic oral conditions including herpes simplex virus, stomatitis, oral lichen planus, angular cheilitis, candidiasis, necrotizing gingival diseases, active systemic infection, active dental caries, or any periodontal diagnosis other than “health periodontium”, or “localized gingivitis” as defined by the European Federation of Periodontology, 2019.

A sample size of 30 participants (15 per group) was initially targeted to detect a statistically significant difference with a hypothesized effect size of 0.4, using a two-tailed test with 5% significance and 80% power, calculated using OpenEpi® Version 3.01. Following screening, 21 eligible participants were enrolled in the parallel arm randomized clinical trial and assigned to either the intervention group (n = 11; nitrate mouthrinse) or the placebo group (n = 10) using a computer-generated random sequence. The allocation sequence was created using a simple randomization method in Microsoft Excel. To ensure allocation concealment, the randomization sequence was generated by an independent researched who was not involved in participant recruitment or data collection.

All study visits, including baseline and follow-up assessments, were conducted at the Faculty of Dentistry clinic. The trial commenced on August 14, 2024, and concluded on November 27, 2024.

Participants in the intervention group were instructed to swish 15mL of a nitrate-rich mouthrinse containing 7.5g of Bio-Steel ® Sport Beets (NPN:80104749) in 250mL of sterile, distilled water, and then expectorate after 30 seconds, once daily for 14 consecutive days, following their usual nightly oral hygiene routine. Participants in the control group were instructed to swish with 15mL sterile, distilled water for 30 seconds, and then expectorate. Participants were instructed to maintain their typical diets and oral care practices throughout the study period.

##### Data Collection and Analysis

At the baseline visit, participants were screened for eligibility criteria and underwent an oral examination to screen for oral pathologies, caries, and periodontal disease.

The primary outcome variable of oral microbiome was collected as tongue swab samples using the OMNIGENE® ORAL (OMR-120) collection kit. A secondary outcome of salivary nitric oxide concentrations was evaluated using HumanN® nitric oxide indicator strips according to the manufacturer’s protocol. The resulting colorimetric readings were compared against a reference legend and categorized on a 3-point scale: depleted, low, or optimal. Standardized photographs of the strips alongside the reference legend were reviewed by two blinded examiners. In cases of disagreement, a third blinded examiner provided the final assessment. Blood pressure was collected as an exploratory variable using a calibrated manual sphygmomanometer and a standard stethoscope, with participants seated and rested for at least 5 minutes prior to measurement. Three readings were taken at 5-minute intervals, with the average value used for analysis in accordance with American Heart Association guidelines.

Follow-up visits occurred exactly 14 days after the baseline visit and at approximately the same time of day to control for diurnal variation.

Tongue swab samples were submitted for 16S rRNA sequencing to characterize and compare the microbial community structure between groups. DNA extraction and quantification were conducted by Genome Québec. A total of 42 bacterial libraries (16S V3–V5, using primers 515bF–926R) and 42 libraries (16S V1–V3, using primers 27Fb–519R), along with one PCR blank per set, were prepared. A standard bioinformatics pipeline was employed, including analyses of alpha and beta diversity, and differential relative abundance of taxa between the intervention and placebo groups.

## Notes

### Competing Interest Statement

The authors have declared no competing interest.

